# Local controls modify the effects of timber harvesting on surface soil carbon and nitrogen dynamics

**DOI:** 10.1101/2024.06.10.598062

**Authors:** Elisabeth B. Ward, Mark S. Ashton, Jessica L. Wikle, Marlyse Duguid, Mark A. Bradford

**Affiliations:** Department of Environmental Science and Forestry, The Connecticut Agricultural Experiment Station, New Haven, CT 06511, USA; The Forest School, Yale School of the Environment, Yale University, New Haven, CT 06511, USA; Rubenstein School of Environment and Natural Resources, University of Vermont, Burlington, VT, 05405, USA

**Keywords:** Downed woody material, legacy tree retention, microbial biomass, regeneration, soil carbon, soil nitrogen

## Abstract

Managing for structural complexity to enhance forest ecosystem health and resiliency is increasingly incorporated in silvicultural treatments. High spatial variability in stands managed for structural complexity could obscure the effects of forest management on surface soils. Yet few studies have assessed how within-stand variation in forest structure and other local controls influence surface soil organic matter dynamics over time following timber harvests. We used a stratified random sampling design to capture variation in stand age, legacy structure, soil type, and topography in a second-growth, oak-hardwood forest in the northeastern U.S. We compared surface soil carbon and nitrogen content and availability in 15 harvested stands managed to promote tree regeneration (*n* = 144 plots) and five unharvested controls (*n* = 48 plots). We also examined changes over time since harvest in just the harvested stands using a 22-year chronosequence. Forest management strongly influenced surface soil carbon and nitrogen dynamics. The timber harvests had lower soil carbon and nitrogen, microbial biomass, and carbon mineralization but higher nitrogen mineralization. These differences were more pronounced in the drier, less fertile soil type than in more moist, fertile soils. Across the 22-year chronosequence, topography, soil type, and downed woody material density dictated the direction of changes in surface soil carbon and nitrogen over time. Soil carbon and nitrogen accrued over time at drier, higher elevation (∼300 m) sites and under higher densities of fine woody material but declined at lower elevations (∼180 m) and under lower fine woody material. Proximity to legacy trees was associated with higher soil carbon and nitrogen concentrations and availability. Our findings underscore the importance of silvicultural practices that retain structural legacies and downed woody material in shaping surface soil carbon and nitrogen dynamics over time. Our results also highlight how accounting for spatial variation in local controls on soil carbon and nitrogen, such as topography, can improve detection of changes from forest management practices that increase spatial heterogeneity within stands, such as irregular shelterwood and seed tree regeneration methods.

## Introduction

Understanding how forest management practices influence soil physical and biological properties and processes is critical for sustaining the long-term health and productivity of forests and the products and services they provide. Globally, forests support most biodiversity on Earth and are the largest, land-based natural climate mitigation opportunity (Griscom et al., 2017; IPCC, 2023). Simultaneously, over 30% of global forests are actively managed for timber production, and billions of people depend on forest products and services for their livelihoods (FAO, 2022). Past intensive forestry practices have, however, often simplified forest structure and composition, which has reduced biodiversity in many forests worldwide (Puettmann et al., 2012). Forestry practices that balance multiple management goals are therefore key to sustaining long-term forest ecosystem health and resiliency while also supporting societal needs through natural resource production (Gustafsson et al., 2012; Hua et al., 2022; IPBES, 2019).

In native, second-growth forests, silvicultural methods to promote natural regeneration and sustain forest ecosystem functioning often enhance forest complexity by emulating natural disturbance regimes that create structural legacies (D’Amato et al., 2017; Long, 2009). In eastern North America, for instance, episodic, catastrophic disturbances include hurricanes and other severe windstorms that result in partial stand mortality and increase downed woody material. Shelterwood and seed tree regeneration methods emulate this natural disturbance regime by creating large canopy gaps interspersed with mature overstory trees to promote the establishment and/or release of advance tree seedling reproduction (Ashton and Kelty, 2018). In uniform shelterwood and seed tree methods, legacy overstory trees are removed once regeneration is secured. In irregular treatments, however, legacy trees are retained in the long term as reserves to enhance the structural complexity of the stand (Raymond et al., 2009). In addition to supporting wildlife (Fedrowitz et al., 2014; Hanle et al., 2020), legacy trees maintain living plant carbon (C) inputs that support belowground communities (Prescott and Grayston, 2023), including mycorrhizal fungi (Kim et al., 2021; Kranabetter et al., 2013; Sterkenburg et al., 2019), and could therefore contribute to soil organic C and nitrogen (N) recovery over time. Downed woody material also contributes to structural complexity and can help enhance soil organic matter (SOM) recovery (Achat et al., 2015; James et al., 2021; Klockow et al., 2013). Yet despite the recent interest in silvicultural practices to enhance structural complexity within stands, few studies have assessed how variation in legacy tree retention and downed woody material influence the effects of timber harvesting on surface soil C and N over time (Kurth et al., 2020).

Changes in soil C and N availability in response to forest management can shape long-term patterns of forest development and productivity (Powers et al., 2005; Schoenholtz et al., 2000; Thiffault et al., 2011), and surface soils are often more vulnerable to disturbances than deeper horizons (James and Harrison, 2016; Mayer et al., 2024; Nave et al., 2010). In moist, temperate forests, microbial processing of SOM is constrained by N availability, resulting in the accumulation of partially decomposed organic material on the forest floor (Currie, 1999; Ponge, 2003). Fine roots, mycorrhizal fungi, and free-living microbes are all concentrated in these surface, organic-rich horizons (∼0-10 cm; Oa and Ah horizons) (Carteron et al., 2021; Wallander et al., 2004; Yanai et al., 2008), where they contribute to plant growth and productivity by enhancing N mineralization (Aber et al., 1993) and improving soil porosity and aggregate formation (Henderson, 1995). However, surface SOM accumulation can also restrict tree seedling germination and growth by increasing soil temperatures and limiting N and water availability (Caspersen and Saprunoff, 2005; Prescott et al., 2000). Disturbances that expose mineral soils and enhance N availability can therefore help promote regeneration (Hupperts et al., 2020; Willis et al., 2015; Zaczek, 2002). Yet over time, recovery of surface soil C and N and microbial biomass helps sustain plant growth and productivity by enhancing soil aeration, water holding capacity, and nutrient availability (Prescott et al., 2000; Thiffault et al., 2011).

Quantifying changes in surface soil C and N in response to forest management is notoriously challenging owing to high fine-scale spatial variation in soil conditions, which can preclude detection of treatment effects (Conant et al., 2003; Homann et al., 2001; Oliver et al., 2004; Yanai et al., 2003b). Past work analyzing soil C and N following timber harvests have reported negligible or idiosyncratic impacts that may simply reflect high spatial variation within stands as opposed to true effects (Carpenter et al., 2021; Nave et al., 2010; Warren and Ashton, 2014). Soil processes are driven by multiple controls that operate at different scales, and the difficulty of reconciling scale-dependent changes can further obscure forest management effects (Bernhardt et al., 2017; Bradford et al., 2021; O’Rourke et al., 2015). Among forest stands, differences in land-use history, topography, soil type, and vegetation age and composition strongly influence surface soil C and N availability (Côté et al., 2000; Jobbagy and Jackson, 2000; Kurth et al., 2014; Waring et al., 2016). Within forest stands, fine-scale biotic and abiotic factors that regulate microbial activity, such as soil moisture, temperature, pH, and plant C inputs, control C and N mineralization rates (Schlesinger, 1997). Accounting for spatial variation in controls on surface soil C and N dynamics could help disentangle the effects of forest management from other confounding factors and reveal contingent outcomes. Soil C and N concentrations also respond slowly to changes in management (Yanai et al., 2003b), so more responsive microbial variables, such as C and N transformation rates, could provide greater insight into trajectories of changes over time in response to silvicultural treatments (Berhe, 2019).

Here, we assess how variation in local controls on soil C and N content and availability mediate the effects of forest management on surface soils over time. We used a stratified random sampling design to capture variation in legacy overstory structure, soil type, and topography within and among forest stands. Specifically, we ask: (Q1) How do surface soil C and N dynamics differ in 15 harvested stands managed to promote tree regeneration relative to five unharvested stands? and (Q2) How do local controls influence changes in surface soil C and N over time since harvest in a 22-year chronosequence? We hypothesized that timber harvesting would reduce surface soil C by decreasing live plant C inputs and through physical disturbances to the organic horizon that enhance SOM decomposition. We also hypothesized that timber harvesting would increase N mineralization due to reduced plant N demand and higher SOM turnover. We expected these effects to vary with soil type, topography, and legacy structure. As such, stratification of sampling by these controls could improve detection of changes in surface soil C and N over time and provide insight into the mechanisms driving these trends.

## Materials and methods

### Study area

We conducted our work at Yale-Myers Forest (41° 57’ N, 72°07’ W), a 3,213-ha mixed hardwood, second-growth forest in Connecticut, USA. The climate is moist and temperate, with a mean January temperature of -4.6°C, a mean July temperature of 21.7°C, and a mean annual precipitation of 133 cm (NOAA, 2021). Soils are well-drained inceptisols of glacial till origin (Table S1), and the topography includes rolling ridges and valleys ∼160 to 320 m in elevation underlain by metamorphic bedrock. Dominant tree species include red (*Quercus rubra* L.), black (*Q. velutina* Lam), and white (*Q. alba* L.) oaks; shagbark (*Carya ovata* (Mill.) K. Koch) and mockernut (*C. tomentosa* (Lam.) Nutt.) hickories; red (*Acer rubrum* L.) and sugar (*A. saccharum* Marshall) maples; black birch (*Betula lenta* L.); white ash (*Fraxinus americana* L.); eastern hemlock (*Tsuga canadensis* (L.) Carriere); and white pine (*Pinus strobus* L.).

European colonial land-use history strongly shapes the structure and composition of the Forest. From 1730 to 1850, about two-thirds of the Forest were cleared by colonists for use as pasture, with the remaining land selectively logged as woodlots (Yale Forests, 2021). Following agricultural abandonment of much of the land in 1850, the Forest recovered to old-field pine, which was subsequently cut in the early 1900s. This common management history resulted in the contemporary predominance of 90- to 120-year-old, second-growth, oak-hardwood stands. Beginning in 1990, when the forest became viably reproductively mature, 10- to 20-ha areas were annually regenerated through irregular shelterwood and seed tree regeneration methods to diversify the structure, composition, and age class distribution within and among stands across the Forest (Yale Forests, 2021; Ashton et al., 2015). These annual regeneration treatments make Yale-Myers Forest well-suited for studying the effects of timber harvesting associated with regeneration treatments over time through a chronosequence approach (Carpenter et al., 2021; Hanle et al., 2020; Wikle et al., 2019).

### Silvicultural treatments and harvest operation

To establish the 22-year chronosequence, we selected 15 stands managed through irregular shelterwood/seed tree regeneration methods from 1994 to 2016. These silvicultural treatments both retain legacy overstory trees as a seed source to promote regeneration but vary in spacing and the extent of site preparation, with treatment severity dependent on site-specific conditions. Irregular shelterwood and seed tree regeneration methods differ from uniform treatments in that legacy overstory trees are permanently retained with irregular spacing (Ashton and Kelty, 2018; Raymond et al., 2009). Shelterwood systems have narrower spacing (35–60 m) to provide partial shelter for the establishment and release of species reliant upon advanced seedling reproduction (e.g., *Quercus* spp.) (Ashton and Kelty, 2018). This regeneration treatment is generally applied on drier, less productive sites more conducive to persistence of oak advance reproduction (Frey et al., 2007; Frey and Ashton, 2018; Nowacki et al., 1990) and where narrower spacing is sufficient owing to higher understory light availability (Fladeland et al., 2003). Seed tree systems have wider spacing (60–80 m) to provide a seed source but minimal shelter and are more often applied on moist, productive sites that have greater competition from mesophytic species (Frey et al., 2007) and lower understory light availability (Fladeland et al., 2003). Seed tree systems are therefore often accompanied by more intensive site treatments (e.g., soil scarification, vegetation crushing) to manage competing understory vegetation and favor the establishment and growth of long-lived, shade-intolerant, pioneer species following the disturbance (Ashton and Kelty, 2018). Site treatments that accompany seed tree methods can cause greater impacts on surface soils compared to shelterwood systems (Ashton and Kelty, 2018). In our study system, however, irregular shelterwood and seed tree treatments are, operationally, end-members along a continuum of varying legacy tree spacing with more minimal differences in site treatments. As such, we do not separate the effects of these two treatments (hereafter “regeneration treatments”) since our study design controlled for spacing relative to legacy trees as well as soil type and topography.

The harvest operation for both regeneration treatments consisted of felling the majority of subcanopy trees and distributing the slash across the ground. Canopy tree spacing varied, with 35–60 m typical of the irregular shelterwood stands and 60–80 m typical for the seed tree treatments. All remaining canopy trees were felled, and the merchantable portions of the boles were removed. Non-merchantable residues were cut and scattered in a layer of slash <1 m in height. Advance regeneration was protected during the harvest operation, and no broadcast soil scarification or prescribed burning was used as a site preparation treatment. However, mountain laurel (*Kalmia latifolia* L.) and other understory shrubs were purposefully crushed where present to release advance regeneration (Moser et al., 1996). Harvesting equipment was also used to scarify the soil surface to promote tree regeneration in areas where recalcitrant, clonal understory herbs, such as hay-scented fern (*Sitobolium punctilobulum* (Michx.) Desv.), were present.

### Sampling design

We stratified our sampling by two soil types, nine timber harvest years, four topographic classes (“terrain classes”), and two distances relative to legacy overstory trees, which yielded 144 plots in 15 harvested stands managed through the regeneration treatments (Fig. 1). We also established 48 plots in five unharvested stands following a similar sampling design (Fig. 1).

**Fig. 1.**
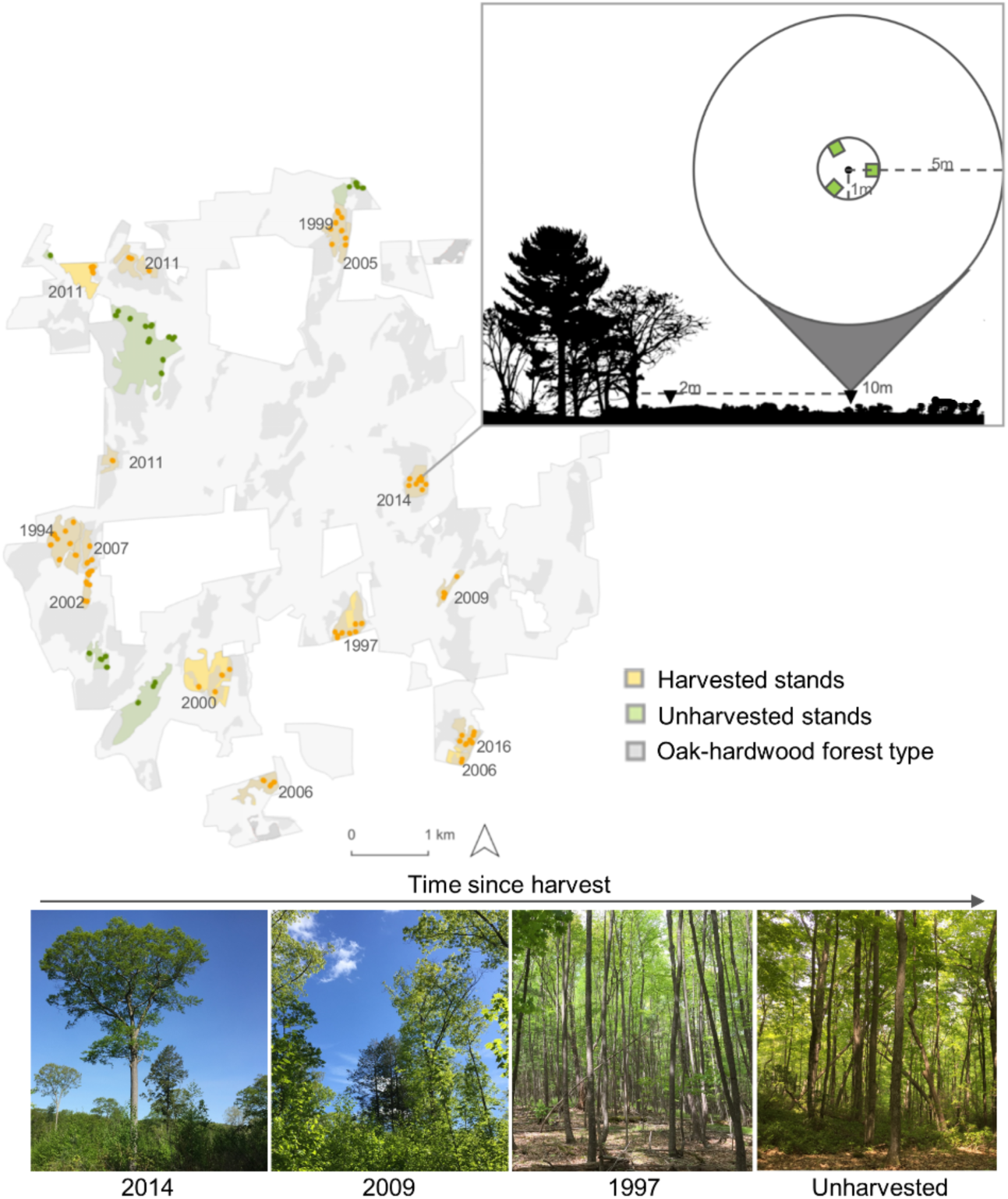
Map and sampling design for the harvested stands managed to promote tree regeneration through irregular shelterwood/seed tree treatments (n = 144 plots) and the unharvested controls (n = 48) at Yale-Myers Forest. Years displayed on the map are the harvest year for the regeneration treatments, and the images show differences in stand structure across the time since harvest chronosequence for different harvest years.

To stratify sampling by soil type, we first identified all U.S. soil series at Yale-Myers Forest (NRCS, 2019; Table S1). We limited our sampling to oak-hardwood stands, which are most commonly managed through shelterwood and seed tree regeneration methods (Ashton and Kelty, 2018; Dey, 2014). Three soil series collectively accounted for 86% of oak-hardwood stands at the Forest: Nipmuck-Brookfield complex (39%), Paxton-Montauk fine sandy loams (26%), and Woodbridge fine sandy loams (21%). We excluded other soil types since they were not adequately replicated (Table S1). We aggregated the Paxton-Montauk and Woodbridge series since they share the same geological features and parent material and differ only in drainage class (well-drained versus moderately well-drained, respectively) (NRCS, 2019). The Nipmuck-Brookfield complex (“ablation tills”) consists of well-drained upland ablation tills derived from schist, granite, and gneiss and tends to be drier, rockier, and less fertile (NRCS, 2019; Fig. S3). The Paxton-Montauk and Woodbridge series (“basal tills”) are mesic, fertile basal tills of drumlins derived from iron-sulfide bearing schist (NRCS, 2019; Fig. S3).

For each of these soil types, we identified areas of stands from nine harvest years that spanned the 1994 to 2016 chronosequence (2 soil types × 9 timber harvest years = 18 study areas) (Fig. 1; Table S2). We preferentially selected stands that included areas with each of the two soil types to capture within-stand heterogeneity. Within these 18 areas, we used the “cluster” function in R (version 4.4.0; R Core Development Team, 2024) with the CLARA K-medoids method (Maechler et al., 2023) to run an unsupervised classification on slope, aspect, and elevation raster bands from publicly available Digital Elevation Model data (CT ECO, 2016) to create four classes with the most distinct topographic features (Fig. S1). We randomly located points at least 20 m from the harvest edge for each of the resulting 72 polygons (2 soil types × 9 timber harvest years × 4 terrain classes). Finally, we established plots 2 m and 10 m from the closest legacy tree (≥20 cm diameter at breast height [DBH] at 1.37 m) to stratify sampling by legacy overstory structure (Fig. 1). This sampling design yielded 144 plots in the harvested stands managed to promote tree regeneration (2 soil types × 9 timber harvest years × 4 terrain classes × 2 distances relative to legacy trees).

For the unharvested stands, we chose five stands with the same soil series (Nipmuck-Brookfield complex and Paxton-Montauk/Woodbridge) that had mean elevation (236–272 m, mean = 251 m) and slope (slope: 1.24–21.1°, mean = 10.7°) values within the range of the plots in the irregular shelterwood/seed tree stands (179–305 m, mean = 246 m; 0.64–28.9°, mean = 9.9°). Across the five stands, we identified three areas with no recorded timber harvests since Yale University acquired the property in 1930, two areas that were crown thinned for firewood 40+ years ago (1973 and 1979), and two areas that were crown thinned for firewood 35-40 years ago (1980 and 1984). We used a similar stratified random sampling design as the regeneration treatments to locate points within each of the unharvested areas (2 soil types × 3 time-since-thinning classes × 4 terrain classes × 2 tree distances = 48 plots). Although we established plots 2 m and 10 m away from the nearest tree ≥20 cm DBH for consistency, each of the paired plots in the unharvested stands had comparable basal area to one another and to the plots 2 m from legacy trees in the harvested stands (Fig. S2).

The contemporary composition of the harvested and unharvested stands was shaped by common indigenous and colonial management histories. All stands originated from colonial old-field pastures, with stands on the more moist, fertile basal till soil series (Paxton-Montauk/Woodbridge) likely from plowed pastures and those on the drier, less fertile ablation till soils (Nipmuck-Brookfield complex) mainly from unimproved pastures. In addition, the unharvested controls were all eligible for the same regeneration treatments as the harvested stands. As such, the five unharvested stands were suitable comparisons for assessing the effects of timber harvesting associated with the regeneration treatments.

### Vegetation and soil sampling

We sampled vegetation and soils in May through June 2019. We identified the species and measured DBH of all living trees and shrubs ≥5-cm DBH in a 5-m radius fixed area plot and ≥1-cm DBH in a nested 1-m radius plot (Fig. 1). We calculated coarse woody material (CWM) volume for the portion of each piece of downed wood >10 cm in diameter that fell within the 1-m radius plot and tallied all fine woody material (FWM) pieces ≥2-cm diameter but less than 10 cm.

We collected and pooled eight 2-cm diameter, 10-cm deep soil cores in the 1-m radius plots. We chose this depth because our focus was not on soil C stocks but rather on the properties and processes occurring in the surface Oa and Ah horizons, which have the highest biological activity in mesic, temperate forest soils. In plots with low soil bulk density, we collected an additional 1–4 cores to ensure we had sufficient sample for the laboratory assays. We recorded the exact depth of each core to calculate bulk density. We kept field collected soils in coolers and stored them at 4°C for less than 4 weeks prior to analysis. We measured volumetric water content (VWC) to a depth of 12 cm at four locations using a HS2 HydroSense Soil Moisture Sensor (Campbell Scientific) and measured the depth of the Oa horizon at three locations within each 1-m radius plot. We also measured soil temperature to a depth of 30 cm using an HI 145 T-Shaped Soil Thermometer (Hanna Instruments, Inc.) and recorded the date and time of day of the measurement for temperature corrections (Methods S1).

### Laboratory assays

We weighed field-moist samples for use in calculating soil bulk density and homogenized and passed each sample through a 4-mm sieve. We mixed samples with water in a 1-to-1 volumetric ratio and measured the pH of the supernatant after 10 min using a benchtop meter (VWR sympHony Sb70p). We measured gravimetric water content (GWC) by oven-drying field-moist samples to constant mass at 105°C. We estimated water holding capacity (WHC) by saturating each soil sample and allowing it to drain freely for 2-h (Paul et al. 2001). We estimated SOM content by calculating mass loss on ignition of soils heated at 550°C for 12-h in a muffle furnace. We used a ball mill to grind air-dried subsamples to a fine powder and analyzed %C and %N with an elemental analyzer (Costech ESC 4010, Costech Analytical Technologies Inc.Valencia, California, USA).

We used a series of laboratory assays to assess microbially available C and N, which we expected be more responsive to management than C and N concentrations. We estimated active microbial biomass with a modified substrate-induced respiration (SIR) method that uses autolyzed yeast extract as a labile C substrate (Fierer et al., 2003; Wardle and Ghani, 1995; West and Sparling, 1986). We then measured rates of CO_2_ efflux over a 4-h incubation period with an Infra-Red Gas Analyzer (Li-COR model Li-7000, Li-Cor Biosciences, Lincoln, Nebraska, USA). To estimate potential C and N mineralization rates, we incubated samples for 30-d at 20°C under a humid atmosphere and maintained 65% WHC, which is within the optimal range for microbial activity (Langenheder & Prosser, 2008; Paul et al., 2001). For the C mineralization assay, we calculated cumulative C-CO_2_ production rates over the 30-d incubation period by integrating CO_2_ efflux measurements sampled over 24-h periods at days 1, 4, 11, 24, and 30 (Fierer et al., 2005). For the 30-d net potential N mineralization assay, we followed protocols modified from Hart et al. (1994) and Robertson (1999). For the initial and day 30 extractions, we shook 6 g dry-weight-equivalent of each soil sample with 25 mL 2M KCl, refrigerated the samples for 12 h, filtered the samples with Whatman Grade 42 filters, and analyzed [NH_4_^+^] and [NO_3_^-^] using a flow analyzer (Astoria 2, Astoria-Pacific, Clackamas, Oregon, USA). Net potential N mineralization was calculated as the difference between the initial and final sum of [NH_4_^+^] and [NO_3_^-^] over the 30-d incubation period.

### Statistical analyses

We used hierarchical linear mixed effects models (LMMs) to analyze differences in surface soil C and N content (%C, %N, and C:N) and availability (C and N mineralization and microbial biomass) between the harvested and unharvested stands (Q1) and over time since harvest in the 22-year chronosequence (Q2). We included plot pair and pair nested within stand as random effects to account for our hierarchical sampling design. We centered all binary variables and centered and standardized all continuous variables by subtracting the mean and dividing by two standard deviations to enable direct comparison of the mean effect sizes of all predictors within the same model (Gelman, 2008). We natural log-transformed microbial biomass to meet assumptions of normal residual distributions. We also natural log-transformed the C:N ratios and interpreted them alongside %C and %N data to minimize inferential issues that arise from analyzing ratio response variables (Isles, 2020; Jasieński and Bazzaz, 1999). We calculated variance explained by fixed (marginal *R*^2^) and by fixed and random (conditional *R*^2^) effects (Nakagawa & Schielzeth, 2013) using the ‘MuMIn’ package (Bartoń, 2023). We fit all models using a Gaussian error distribution and ran all analyses in R (version 4.0.3; R Core Development Team, 2024).

To address Q1, we compared surface C and N content and availability in the harvested stands managed through regeneration treatments (*n* = 144 plots) and the unharvested controls (*n* = 48 plots). Owing to the unbalanced sample sizes and unequal variances, we ran these LMMs using the ‘lme’ function from the ‘nlme’ package, which can be used to model heteroscedasticity for different levels of a group (Pinheiro and Bates, 2000). We specified unequal variances within groups using the ‘weights’ argument with ‘varIdent.’ Fixed effects in these models included timber harvesting (binary; harvested = 1, unharvested = 0), proximity to legacy tree (binary; 2 m = 1, 10 m = 0), soil type (binary; ablation tills = 1, basal tills = 0), and the two-way interactions between timber harvesting and the other predictors. We did not include elevation as a fixed effect in these LMMs owing to the wider ranges in elevation in the harvested (179–305 m) versus unharvested (236–272 m) plots.

To address Q2, we ran a second set of LMMs for the plots in the harvested stands only using the ‘lmer’ function in the ‘lme4’ package (Bates et al., 2015). Fixed effects in these models included time since harvest (years; continuous), proximity to legacy tree (binary; 2 m = 1, 10 m = 0), soil type (binary; ablation tills = 1, basal tills = 0), elevation (continuous), FWM density (continuous), and all two-way interactions between time since harvest and the other predictors. We included FWM density in these models owing to the potentially strong effects of downed woody material on soil C and N (e.g., Achat et al., 2015). We focused on FWM and not CWM since our random sampling included a high number of plots with no CWM, which precluded meaningful interpretation of the effects of CWM. Our original sampling design did not include stratification by downed woody material since we had anticipated its density to covary with time since harvest. Such mediating variables steal variance and hence reduce the effect size of the primary cause (e.g., harvest) when included in the same statistical models (McElreath, 2016). However, we found no evidence for a relationship between FWM density and the other focal predictors. The square-root of the variance inflation factors for the main effects were also less than 2, indicating low collinearity amongst the predictor variables.

Given that we expected many of our measured soil variables to mediate the effects of the regeneration treatment on surface soil C and N, we used principal component analysis (PCA) to visualize multivariate trends between all measured soil variables in the harvested versus unharvested plots (Q1) and over time since harvest in the 22-year chronosequence (Q2) (Fig. 2). We natural log transformed GWC, WHC, %SOM, net potential N mineralization, and microbial biomass to increase the linearity of the relationships among the soil variables, which improves PCA performance (Legendre and Legendre, 2012). We excluded one harvested plot with a missing WHC value. We tested the significance of relationships between the regeneration treatment and time since harvest and the scores of the first two principal components using simple linear regression. We ran all PCAs using the ‘princomp’ function in R.

**Fig. 2.**
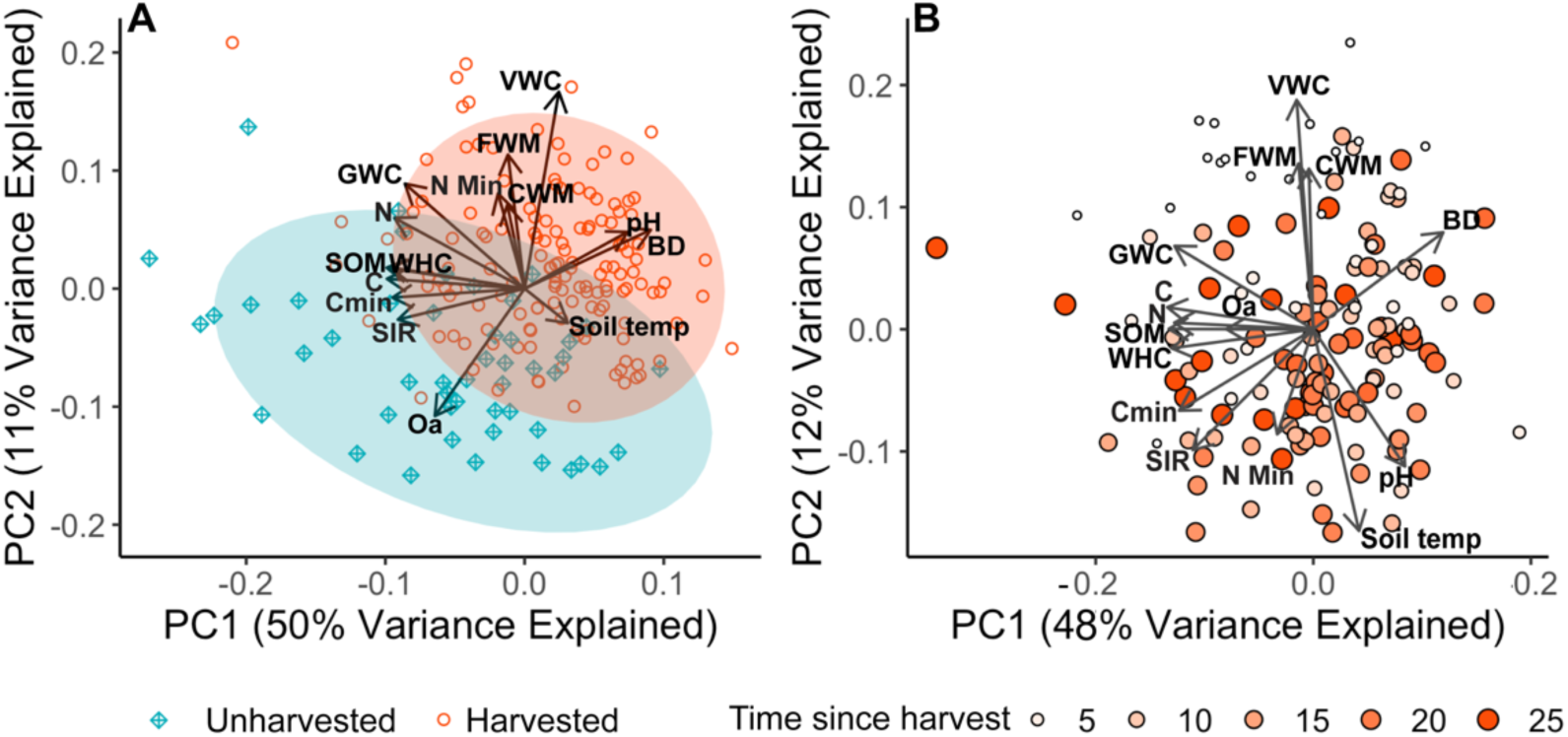
Principal component analysis illustrating multivariate differences in all measured soil variables between (Q1) (A) the harvested and unharvested stands and (Q2) (B) over time since harvest in the 22-year chronosequence. Abbreviations are as follows: volumetric water content (VWC), fine woody material (FWM), coarse woody material (CWM), net potential nitrogen mineralization (N Min), bulk density (BD), soil temperature (Soil temp), average Oa horizon depth (Oa), active microbial biomass as measured through substrate induced respiration (SIR), carbon mineralization (Cmin), carbon content (C), nitrogen content (N), soil organic matter content (SOM), water holding capacity (WHC), and gravimetric water content (GWC).

## Results

### Effects of timber harvesting in the harvested versus unharvested stands (Q1)

Relative to the unharvested controls, the harvested stands had lower soil C and N content and C availability and higher net potential N mineralization rates (Table 1; Figs. 2-5; Tables S3-S4). Mean %C and %N were ∼40% and ∼20% lower in the harvested stands managed through regeneration treatments (*n* = 144 plots) compared to the unharvested controls (*n* = 48 plots). Because there was a stronger negative effect of the regeneration treatments on %C than %N, we also observed a decrease in soil C:N in the harvested stands (Fig. 3; Table S3). Both microbial indicators of surface soil C availability were also lower in the harvested stands compared to the unharvested controls (Figs. 4-5; Table S4). Plots in harvested stands managed to promote regeneration had 27% lower mean C mineralization rates and 58% lower mean microbial biomass compared to the unharvested plots (Table 1). Net potential N mineralization, by contrast, was 39% higher in the regeneration treatments (Table 1; Figs. 4-5), though this difference was not statistically significant.

**Fig. 3.**
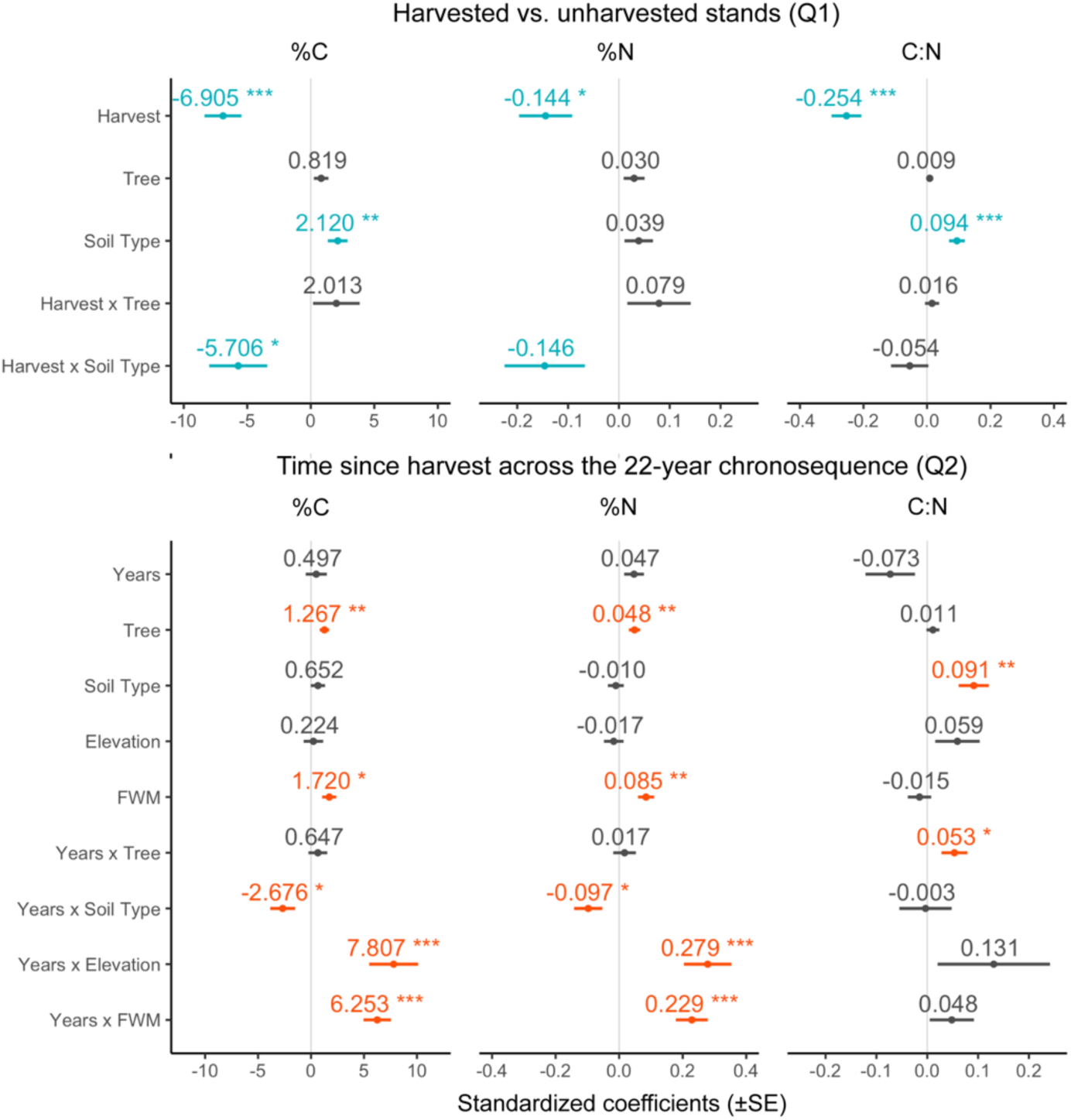
Standardized coefficients of models assessing surface soil carbon (C) and nitrogen (N) content in the (Q1) harvested stands managed through regeneration treatments versus unharvested controls and (Q2) over time since harvest in the 22-year chronosequence. Colors indicate significant (P < 0.05) and marginally significant (P < 0.1) coefficients for the harvested versus unharvested (Q1) (blue) and time since harvest (Q2) (orange) models.

**Fig. 4.**
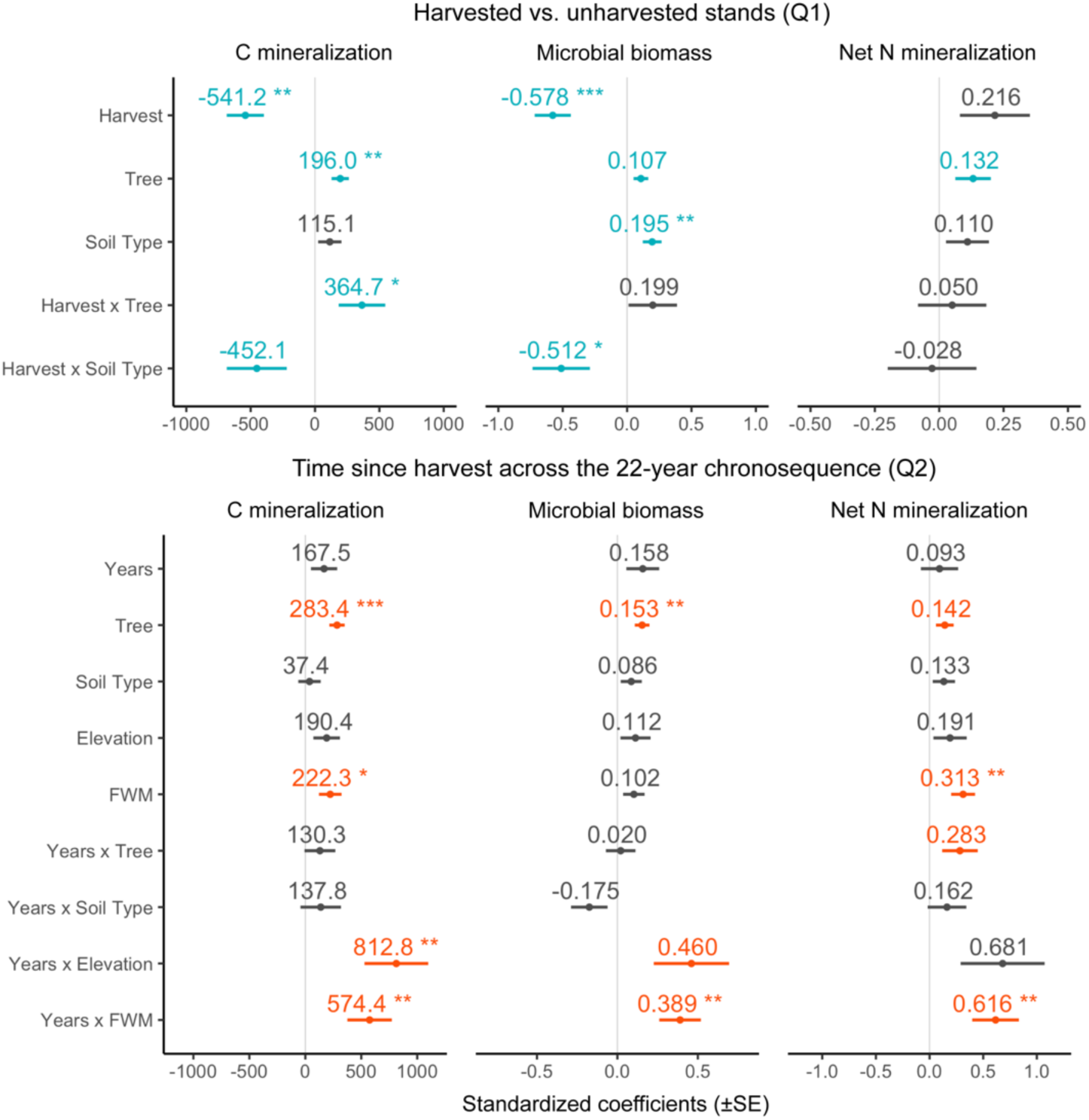
Standardized coefficients of models assessing surface soil carbon (C) and nitrogen (N) mineralization rates and microbial biomass in (Q1) the harvested stands managed through regeneration treatments versus unharvested controls and (Q2) over time since harvest in the 22-year chronosequence. Colors indicate significant (P < 0.05) and marginally significant (P < 0.1) coefficients for the irregular shelterwood versus unharvested (Q1) (blue) and irregular shelterwood chronosequence (Q2) (orange) models.

**Fig. 5.**
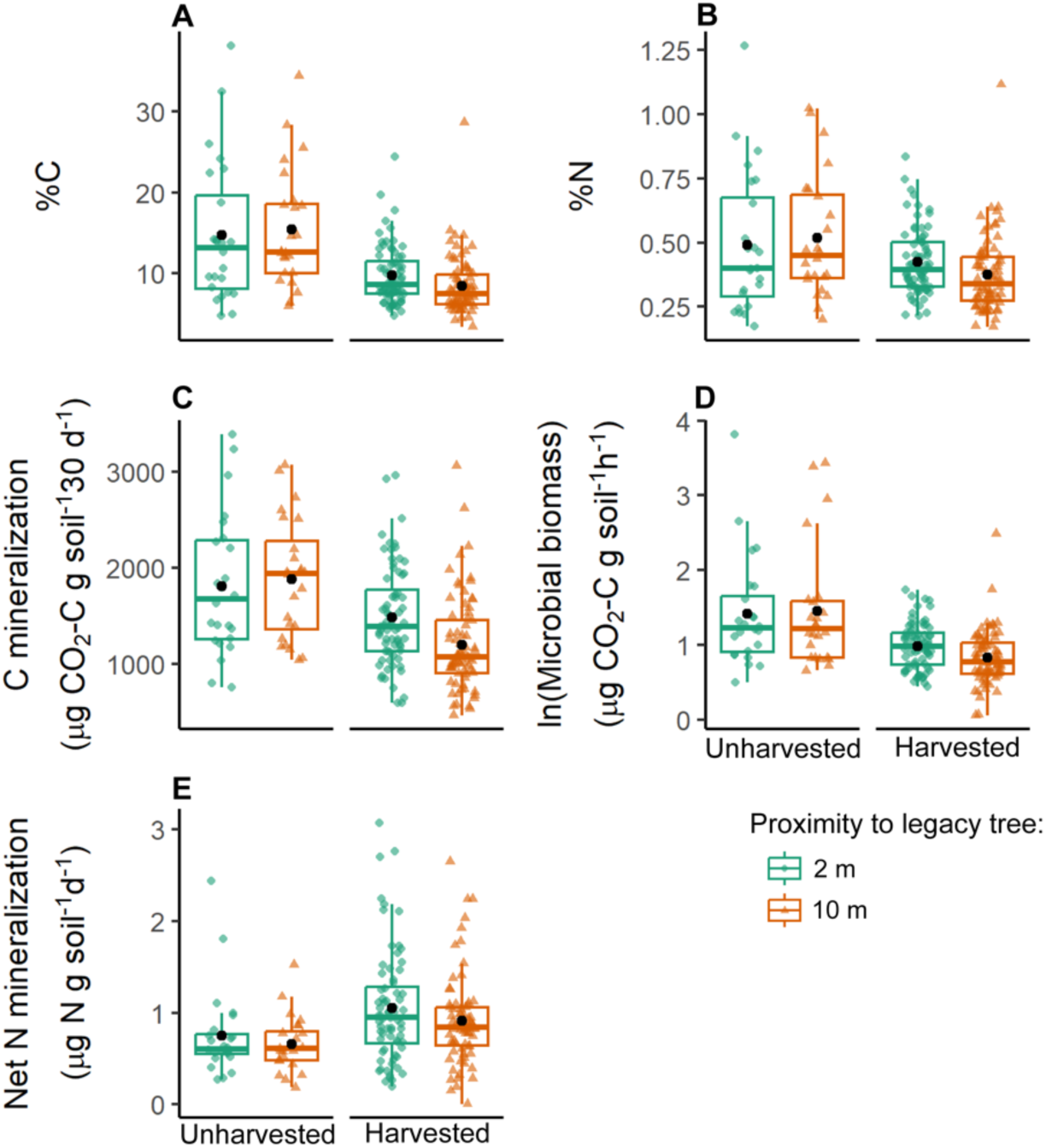
The effects of timber harvesting and proximity to overstory trees on surface soil carbon (C) and nitrogen (N) content (A, B) and availability (C-E). The timber harvests associated with the regeneration treatments negatively influenced surface soil C and N, but proximity to legacy overstory trees (2 m versus 10 m) was associated with higher C and N content and availability. For C mineralization, legacy trees partially mitigated the negative effects of the timber harvest. Black points show the mean values, boxplots show the median, first, and third quartiles, and whiskers show the upper (1.5 × interquartile range) and lower range (1.5 × interquartile range), excluding outliers.

**Table 1.** Surface soil properties and processes in the harvested stands managed to promote tree regeneration through irregular shelterwood/seed tree regeneration treatments (n = 144 plots) and in the unharvested controls (n = 48 plots).

| Soil Variable | Harvested stands |  | Unharvested stands |  |
| --- | --- | --- | --- | --- |
|  | Mean | SD | Mean | SD |
| *Soil microclimate conditions* |  |  |  |  |
| Coarse woody material ($\text{m}^3 \text{ha}^{-1}$) | 124 | 423 | 70.8 | 145.8 |
| Fine woody material (pieces $\text{m}^{-2}$) | 1.90 | 2.00 | 1.46 | 0.90 |
| Oa horizon depth (cm) | 2.05 | 1.32 | 3.83 | 1.51 |
| Soil temperature ($^{\circ}\text{C}$) | 10.25 | 1.26 | 9.85 | 0.75 |
| Mean volumetric water content (VWC) (%) | 28.00 | 7.22 | 17.63 | 6.42 |
| Gravimetric water content (GWC) (%) | 37.11 | 6.95 | 38.26 | 8.64 |
| *Soil physical and chemical properties* |  |  |  |  |
| Bulk density (BD) ($\text{g cm}^{-3}$) | 0.40 | 0.12 | 0.24 | 0.09 |
| Water holding capacity (WHC) (%) | 50.51 | 7.45 | 56.38 | 9.68 |
| pH | 5.02 | 0.35 | 4.49 | 0.30 |
| %SOM | 15.77 | 6.17 | 23.06 | 10.89 |
| %C | 9.05 | 3.69 | 15.05 | 8.04 |
| %N | 0.40 | 0.14 | 0.50 | 0.25 |
| C:N | 22.63 | 3.05 | 29.68 | 3.93 |
| *Soil carbon and nitrogen processes* |  |  |  |  |
| C mineralization<br>($\mu\text{g CO}_2\text{-C g soil}^{-1} 30 \text{ d}^{-1}$) | 1344.3 | 522.7 | 1845.7 | 671.4 |
| Active microbial biomass<br>($\mu\text{g CO}_2\text{-C g soil}^{-1} \text{h}^{-1}$) | 2.64 | 1.18 | 6.34 | 8.51 |
| Net potential N mineralization<br>(Net N Min) ($\mu\text{g N g soil}^{-1} \text{d}^{-1}$) | 0.98 | 0.56 | 0.71 | 0.40 |

Proximity to legacy overstory trees was associated with higher C and N availability (Figs. 3-5; Table S4). Mean microbial biomass and C and N mineralization for all plots in both the regeneration treatments and the unharvested controls were 3, 14, and 16% higher 2 m from legacy trees compared to 10 m. For C mineralization, proximity to legacy trees partially diminished the negative effect of the regeneration treatment (Figs. 4-5; Table S4). The stands managed through regeneration treatments had 36% lower C mineralization in plots 10 m from legacy overstory trees but only 18% lower C mineralization in plots 2 m from overstory trees. We observed similar, moderating effects of proximity to legacy trees in the harvested stands managed through regeneration treatments for %C, %N, and microbial biomass as indicated by the positive coefficients for the Harvest by Tree interaction terms, but these effects were not statistically significant (Figs. 3-5; Table S4).

The effects of timber harvesting associated with the regeneration treatments also varied by soil type (Figs. 3-4, S4; Tables S3-S4). In the unharvested stands, C mineralization and microbial biomass were 32 and 190% higher, respectively, in the drier, less fertile ablation tills compared to the more moist and fertile basal tills. Likewise, C and N concentrations were 68 and 45% higher in the ablation tills than in the basal tills in the unharvested stands. However, differences in surface C and N content and C availability between the soil types were minimal in the irregular shelterwood/seed tree plots, indicating that timber harvesting consistently had stronger negative effects on surface soil C and N concentrations and C availability in the less fertile ablation tills than in the basal tills (Figs. 3-4, S4; Tables S3-S4).

We also observed multivariate differences between all measured soil variables in the harvested and unharvested stands (Table 1, Fig. 2). Compared to the unharvested stands, the regeneration treatment was positively associated with PC1 (Estimate ± SE = 2.78 ± 0.41; *P* < 0.0001; Adjusted *R*^2^ = 0.19), which accounted for 50% of variance and was negatively associated with C and N concentrations and C availability and positively associated with bulk density, pH, and soil temperature (Fig. 2; Tables S3-S4). The regeneration treatment was also positively associated with PC2 (Estimate ± SE = 1.50 ± 0.19; *P* < 0.0001; Adjusted *R*^2^ =0.25), which explained an additional 11% of variance and was positively associated with soil moisture, N mineralization, and downed woody material density and negatively associated with the depth of the Oa horizon (Fig. 2; Tables S3-S4).

### Changes over time since harvest in the 22-year chronosequence (Q2)

We observed high within and among stand variation in surface soil C and N content and availability over time since harvest across the chronosequence (Figs. 6, S5). Variation among plots within stands of the same age typically exceeded variation in the mean values by stand age across the chronosequence. For instance, the standard deviation of the mean %C values for each stand age was 1.4%, but the standard deviations for %C ranged from 1.8 to 5.8% within stands of the same age.

**Fig. 6.**
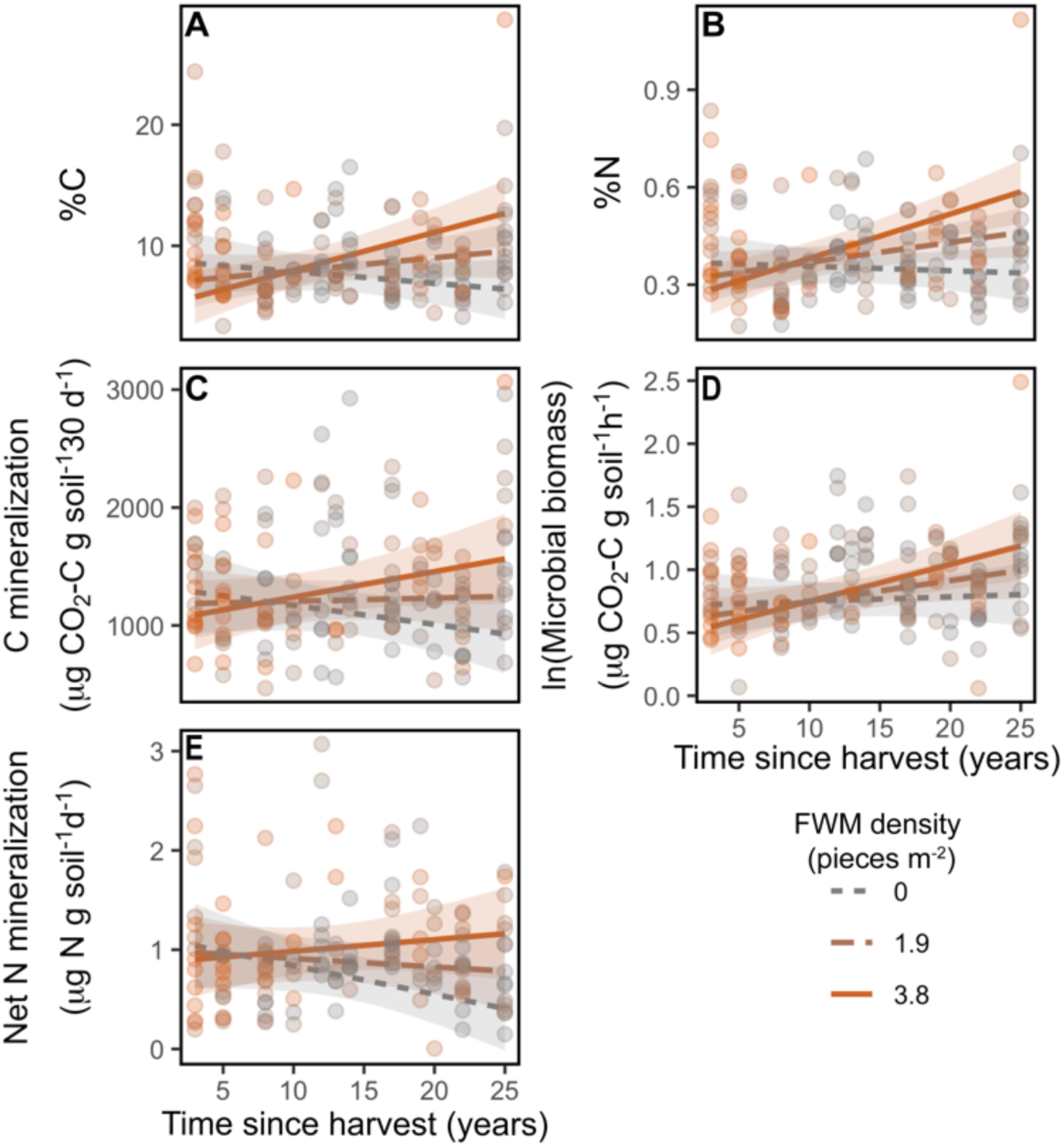
The recovery of surface soil carbon (C) and nitrogen (N) content and availability over time since harvest (n = 144) was contingent on the presence of fine woody material (FWM). The regression lines depict the modeled effects of C and N content and availability from the linear mixed effects models (Tables S5-S6), which include plot pair and pair nested within stand as random effects.

Despite this high within-stand variation, we observed marked changes in surface C and N content and availability over time. However, these changes were strongly contingent on factors that varied within and among stands. Differences in elevation and FWM density modified the direction and magnitude of time since harvest on surface soil C and N content and availability (Figs. 3-4, 6, S5; Tables S5-S6). Soil C and N increased over time at higher elevation sites and in plots with higher FWM density but declined or remained constant in lower elevation sites and in plots with lower FWM density (Figs. 3-4, 6-7; Tables S5-S6). Soil type also modified the effects of time since harvest on soil C and N concentrations (Fig. 3; Table S5). Carbon and N concentrations decreased slightly over time in the drier, less fertile ablation tills but increased in the more moist and fertile basal tills. However, the strength of this interaction was weak compared to that of FWM and elevation (Fig. 3; Table S5).

Proximity to legacy trees within the harvested stands in the 22-year chronosequence was positively associated with surface soil C and N content and availability (Figs. 3-4, S6; Tables S5-S6). Mean %C and %N were 16 and 13% higher in plots 2 m from legacy trees than in the plots 10 m from legacy trees, and mean C mineralization, microbial biomass, and net N mineralization rates were 24, 13, and 16% higher in plots 2 m from legacy trees. For the most part, this effect was invariant over time since harvest (Figs. 3-4, Fig. S6; Tables S5-S6). However, N mineralization increased over time in plots 2 m from legacy trees but decreased in plots 10 m from trees (Fig. S6; Table S6). Proximity to legacy trees was also associated with a less negative effect of time since harvest on soil C:N, but this effect was small (Fig. 3; Table S5).

We observed high variation in the PCA of surface soil conditions for plots in the harvested stands managed through regeneration treatments, particularly for variables associated with C content and availability (Fig. 2b). Principal component 1 explained 48% of variance and was negatively associated with C and N content and C availability and positively associated with soil bulk density and pH. However, time since harvest was not significantly associated with PC1. Instead, time since harvest was only negatively associated with PC2 (Estimate ± SE = -0.067 ± 0.014; *P* < 0.0001; Adjusted *R*^2^ = 0.13), which explained 12% of variance and was negatively associated with soil temperature, pH, and N mineralization and positively associated with soil moisture and downed woody material (Table S4).

## Discussion

Management practices to enhance structural complexity are increasingly incorporated in silvicultural treatments to help balance multiple management goals, including timber production, natural regeneration, and ecological functions and services (D’Amato et al., 2017; Littlefield & Keeton, 2012; Long, 2009). These practices include the retention of legacy overstory trees and downed woody material (Klockow et al., 2013). Yet enhancing structural complexity could also increase heterogeneity in surface soil C and N (Kurth et al., 2020), which already exhibit high spatial variation among microsites and across gradients of soil type and topography. Structural complexity within stands could therefore obscure forest management effects on surface soils, particularly when sampling densities are low, randomized, and/or aggregated across whole stands (Yanai et al., 2003b). Although higher density sampling at the stand scale could provide robust estimates of the mean harvest effect size for many stands, it is unlikely to resolve mean changes within a single stand (Bradford et al., 2023). Moreover, these study designs can conceal the mechanisms underlying observed changes (Kurth et al., 2020), which have important implications for the design of silvicultural prescriptions. Our study therefore investigates how local variation in topography, soil type, legacy tree retention, and downed woody material influence the effects of timber harvesting on surface soil C and N over time.

### Timber harvesting was associated with lower surface soil carbon and legacy trees enhanced soil carbon availability

In line with our hypotheses, we found lower surface soil C and N content and C availability but higher net N mineralization rates in the harvested stands managed through regeneration treatments compared to the unharvested controls. Physical disturbances from harvesting can fragment, mix, and compact the surface organic horizons, thereby exposing mineral soils and increasing soil moisture (Mayer et al., 2020). We found evidence for each of these physical changes and increased soil temperature (Table 1, Fig. 2a) owing to higher solar radiation on the forest floor in the canopy gaps created to promote regeneration. These changes all enhance organic matter decomposition rates (Covington, 1981; Mayer et al., 2017), which, together with reduced C fixation and live plant C inputs, can decrease soil C content and availability, especially in the surface mineral and organic horizons (Carpenter et al., 2021; James and Harrison, 2016; Mayer et al., 2024; Nave et al., 2010).

We also found that the effects of timber harvesting on surface soil C and N varied by soil type. Differences in C and N content and C availability in the harvested versus unharvested plots were more pronounced in the drier ablation till soils compared to the more moist and fertile basal till soils. Colonial land-use legacies associated with differences in soil moisture and fertility between these soil types could help explain these trends. Whereas the fertile, basal tills were plowed and maintained for a longer period of time as improved pastures, the less fertile ablation till soils were managed less intensively and for a shorter period of time. Compared to unimproved pastures, historically tilled forest soils have lower C and higher N availability (Bárcena et al., 2014; Guo and Gifford, 2002; Laganière et al., 2010), and these effects can persist long-term as soil legacies in second-growth forests in the region (Compton and Boone, 2000). However, these land-use legacies also covary with natural gradients in soil fertility, making it difficult to disentangle their relative influence.

The greater accumulation of more labile, particulate SOM in the Oa horizon on the less fertile ablation tills (Ponge, 2003) could also make these C and N pools more sensitive to losses from disturbances, such as timber harvests (Lavallee et al., 2020). Higher microbial activity and faster SOM turnover on the more moist, fertile basal tills, by contrast, could generate more stable organic C and N in mineral-rich surface horizons (Lavallee et al., 2020; Prietzel et al., 2020). Such mechanisms would be consistent with global meta-analyses linking sites with higher initial surface soil C concentrations, such as N-limited forests with thick organic horizons, to greater susceptibility to soil C losses from disturbances (Crowther et al., 2016; García-Palacios et al., 2024; Mayer et al., 2024). Regardless of the specific mechanisms, our data indicate that surface soil C on drier, less fertile ablation tills may be more susceptible to disturbances from timber harvesting than the more mesic basal till soils.

Because surface SOM accumulation can restrict tree seedling germination and growth by limiting soil moisture and N availability (Caspersen and Saprunoff, 2005; Ponge, 2003; Prescott et al., 2000), lower surface soil C in the harvested stands could, theoretically, enhance tree regeneration. Indeed, we observed slightly higher N mineralization rates in the stands managed through regeneration treatments compared to the unharvested stands. Reduced plant N demand, higher soil temperatures, greater soil moisture availability, and more rapid SOM turnover can all enhance soil N availability following harvests (Piatek and Allen, 1999; Vitousek and Matson, 1985; Waide et al., 1988). These results are consistent with past work showing higher plant available N, soil moisture, and temperatures in canopy gaps compared to closed-canopy stands (Devine and Harrington, 2008; Hawkins and Moran, 2003; Walters et al., 2006). While our study did not address how changes in surface soil C and N dynamics influence tree regeneration following harvests, our observations of higher soil moisture and N availability suggest that disturbances from harvesting may make soil conditions more suitable for regenerating trees (Martin et al., 2021). At broad scales, there can be an inverse relationship between belowground C and aboveground forest productivity (de la Cruz-Amo et al., 2020; Terrer et al., 2021), and there may be a similar trade-off for management practices that promote surface soil C versus tree regeneration.

Proximity to overstory trees was also associated with higher surface soil C within the regeneration treatments, and this effect did not vary with time since harvest. Retention of legacy trees could therefore help maintain above- and belowground living plant C inputs that can support belowground communities and ecological functions (Prescott and Grayston, 2023). Previous work has similarly shown that partial harvests can help mitigate the negative effects of forest management activities on surface soil C compared to more intensive harvests (Simard et al., 2020; Zhou et al., 2013). In a review of the effects of legacy tree retention on soil organic C, Prescott & Grayston (2023) recommend distributing legacy trees throughout harvested stands and maintaining tree distances less than 15 m to maximize benefits to the soil community. Yet, spacing of legacy trees also affects the composition and abundance of regenerating trees by selecting for species with different shade tolerances. Another project conducted in the same stands as our study found that higher legacy overstory basal area reduced the competitive advantage of regenerating oak trees in favor of more shade-tolerant species (Wikle et al., 2019). Legacy tree retention to support belowground communities and promote soil C and N recovery must therefore consider trade-offs with conditions that promote desired tree regeneration. The irregular spacing and arrangement of residual trees and reserves within the regeneration treatments could help balance these opposing objectives by creating heterogeneous soil conditions that better support different management goals across different areas of the stand.

### Detecting changes in surface soil carbon and nitrogen over time following timber harvests

The recovery of surface soil C and N content and availability over time since harvest in the 22-chronosequence was strongly contingent on factors that varied within and among stands. Changes in surface soil C and N depended on topographic position, downed woody material density and, to a lesser extent, soil type. Surface soil C and N accrued with time since harvest at drier, higher elevation (∼300 m) sites and under higher densities of downed woody material but declined at lower elevations (∼180 m) and under lower downed woody material densities. The recovery of surface soil C and N over time at higher topographic positions could be caused by conditions that slow microbial activity and promote SOM accumulation in our study system, such as lower soil moisture (Fig. S3). In the more moist, lower elevation sites, by contrast, rapid decomposition of the pulse of organic residues that immediately follows timber harvests could explain the negative relationship between surface soil C and N and time since harvest. However, the more fertile, basal till soils also recovered C and N over time, whereas soil C and N declined slightly in the more infertile, ablation till soils. These changes were small compared to the effects of elevation and downed woody material but suggest that more work is required to resolve the mechanisms underlying losses or gains in surface soil C and N following timber harvests.

Like elevation, downed woody material also strongly controlled surface soil C recovery over time since harvest across the 22-year chronosequence. Past work investigating the impacts of varying levels of organic matter removal (e.g. whole tree versus bole only removals) on soils and forest productivity have similarly found that downed woody material helps protect and restore surface soil organic C following timber harvests (Achat et al., 2015; James et al., 2021; Klockow et al., 2013; Warren and Ashton, 2014). Fine woody material also helped maintain elevated N availability over time at our study site, whereas N mineralization declined in plots with lower fine woody material densities. Initial immobilization of N by fresh dead wood inputs with high C:N could help retain N in the system, where it can be gradually mineralized over time at more advanced downed wood decay stages (Metzger et al., 2008; Ricker et al., 2016). Although our study design only enabled us to assess the effects of FWM (2-10 cm in diameter) rather than CWM (>10 cm in diameter), we would expect similar effects on surface soil C recovery and N mineralization under CWM but over a longer timeframe owing to the slower decomposition rates and higher C:N of larger pieces of downed wood.

The opposing trajectories of changes in surface soil C and N at different elevations and downed woody material densities illustrate how changes in surface soil C and N can be obscured when sampling is aggregated at the stand scale and local controls are not accounted for through sampling design and statistical modeling. For example, previous work at our study site found reduced surface soil C and N content in the regeneration treatments compared to unharvested stands, which is consistent with our results (Carpenter et al., 2021; Warren and Ashton, 2014). However, they observed no significant changes in C concentrations over the >20 year chronosequence, which contrasts with our work (Carpenter et al., 2021; Warren and Ashton, 2014). One limitation of the chronoseqence approach that could explain these conflicting results is that differences in site conditions among stands can confound or obscure changes over time (Yanai et al., 2003a, 2000). Our results suggest that such issues can, in part, be overcome when sampling designs stratify by controls that vary within and among stands, permitting quantification of temporal changes in surface soil C and N following timber harvests through a chronosequence approach.

## Conclusion

We found that local controls that vary within stands, such as topography and downed woody material, strongly modulated the magnitude and direction of the effects of timber harvesting on surface soil C and N availability across a 22-year chronosequence of stands managed through irregular shelterwood and seed tree regeneration treatments. Recovery of surface soil C and N content and C availability over time only occurred at higher topographic positions and was contingent on the presence of downed woody material. The effects of the regeneration treatments were also moderated by soil type. Compared to the unharvested controls, the harvested stands managed through regeneration treatments had more pronounced differences in surface soil C on the drier, less fertile ablation tills compared to the more mesic and fertile basal tills. Proximity to legacy trees was also associated with higher soil C and N in the harvested stands. These results highlight the importance of retaining structural legacies and downed woody material to recover surface soil C and N over time following silvicultural treatments to promote regeneration. However, these results must be considered alongside potential trade-offs with conditions that favor desired tree species regeneration. Our study also illustrates how accounting for spatial variation in controls on soil C and N, such as topographic gradients and soil type, can improve detection of changes from forest management practices, particularly when they increase spatial heterogeneity within stands by promoting structural complexity.

## Author contributions

E.B.W and M.A.B conceived the ideas and designed the methodology; M.S.A. oversaw the implementation of and maintained records for the irregular shelterwood/seed tree regeneration treatments; E.B.W. collected the data with contributions from J.W. and M.A.B; E.B.W. analyzed the data and led the writing of the manuscript with input from M.A.B. All authors contributed critically to the drafts and gave final approval for publication.

## Acknowledgments

Funding was provided by the Lewis B. Cullman (New York Botanical Garden) and Kohlberg-Donohoe (The Forest School, Yale School of the Environment) Fellowships. Thanks to the Yale Forests faculty, staff, and facilities; the Ingalls Field Ecology Internship Program; the Yale Analytical and Stable Isotope Center; Meghan Midgley; Skylar Albertson; Nicholas Fields; Javier Gonzalez-Rivero; Ki’ila Salas; and David Woodbury.

## Conflict of Interest

The authors declare no conflict of interest.

## Data Availability Statement

Data and code will be available on Dryad Digital Repository upon manuscript acceptance.

## Notes

### Competing Interest Statement

The authors have declared no competing interest.

