## Supplementary Material for "Local controls modify the effects of timber harvesting on surface soil carbon and nitrogen dynamics"

The following Supporting Material is available for this article:

**Methods S1** Soil temperature corrections.

**Fig. S1** Schematic diagram illustrating how we stratified plot locations by topographic variation within each sampling area**.**

**Fig. S2** Variation in tree basal area in plots 2 m and 10m from legacy overstory trees across the (A) time since harvest chronosequence and (B) in the harvested versus unharvested plots.

**Fig. S3** Relationship between elevation, soil type, and soil moisture (volumetric water content) in the harvested plots managed through regeneration treatments ($n$ = 144).

**Fig. S4** The effects of timber harvesting on surface soil carbon and nitrogen content and availability varied by soil type.

**Fig. S5** The recovery of surface soil carbon and nitrogen content and availability over time since harvest in the 22-year chronosequence ($n$ = 144) was contingent on elevation.

**Fig. S6** Influence of proximity to legacy overstory trees on surface soil C and N content across the time since harvest chronosequence.

**Table S1** The frequency and abundance of all U.S. soil series in oak-hardwood forest stands at Yale-Myers Forest.

**Table S2** Harvest years of sampled stands across the 22-year chronosequence for each of the two dominant U.S. soil series used to stratify sampling by variation in geology and parent material.

**Table S3** Unstandardized coefficients of models assessing surface soil carbon (C) and nitrogen (N) concentrations and C:N in the harvested stands versus unharvested controls (Q1).

**Table S4** Unstandardized coefficients of models assessing carbon mineralization, microbial biomass, and nitrogen mineralization in the harvested stands versus unharvested controls (Q1).

**Table S5** Unstandardized coefficients of models assessing surface soil carbon (C) and nitrogen (N) concentrations and C:N in the 22-year chronosequence (Q2).

**Table S6** Unstandardized coefficients of models assessing carbon mineralization, microbial biomass, and nitrogen mineralization in the 22-year chronosequence (Q2).

**Methods S1** Soil temperature corrections.

We used hourly air temperature data from the meteorological station at Yale-Myers Forest to correct the soil temperature measurements for daily and monthly variation, thereby allowing us to analyze these temperature data as though they were collected at the same time and date. Specifically, we standardized and centered the soil and air temperature values to put them on the same scale by subtracting the mean and dividing by the standard deviation. We interpolated the standardized, hourly air temperature values from the meteorological station to get a predicted value at the time of each soil measurement. We subtracted the predicted, standardized air temperature value from the observed, standardized soil temperature value to correct for variation in air temperature. Finally, we back calculated the corrected soil temperature value by adding the mean and multiplying it by the standard deviation of soil temperature.

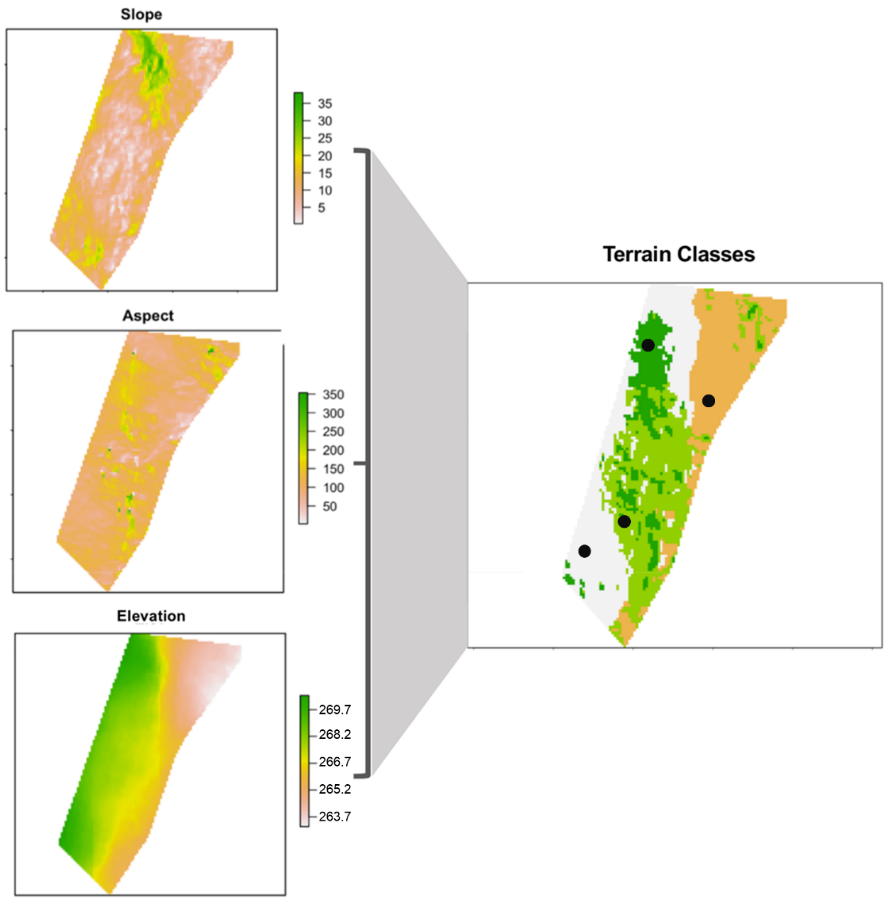


**Fig. S1** Schematic diagram illustrating how we stratified sampling by topographic variation within each of the 18 polygons defined by the combination of two soil types and nine harvest years (Table S2). The figure shows variation in slope, aspect, and elevation within a single such polygon (left side of figure). We then used unsupervised K-medoid classification to create four terrain classes (right side of figure) with the most distinct topographic features based on this raw slope, aspect, and elevation raster data (CT ECO, 2016). The resulting four terrain classes are indicated by the four colors in the right-side panel and have no meaningful interpretation but instead reflect variation in slope, aspect, and elevation within each of the 18 polygons. We randomly located points (black dots) within each of these terrain classes and navigated to these locations in the field, where we established plots 2 m and 10 m from the nearest legacy overstory tree ≥20 cm diameter at breast height.

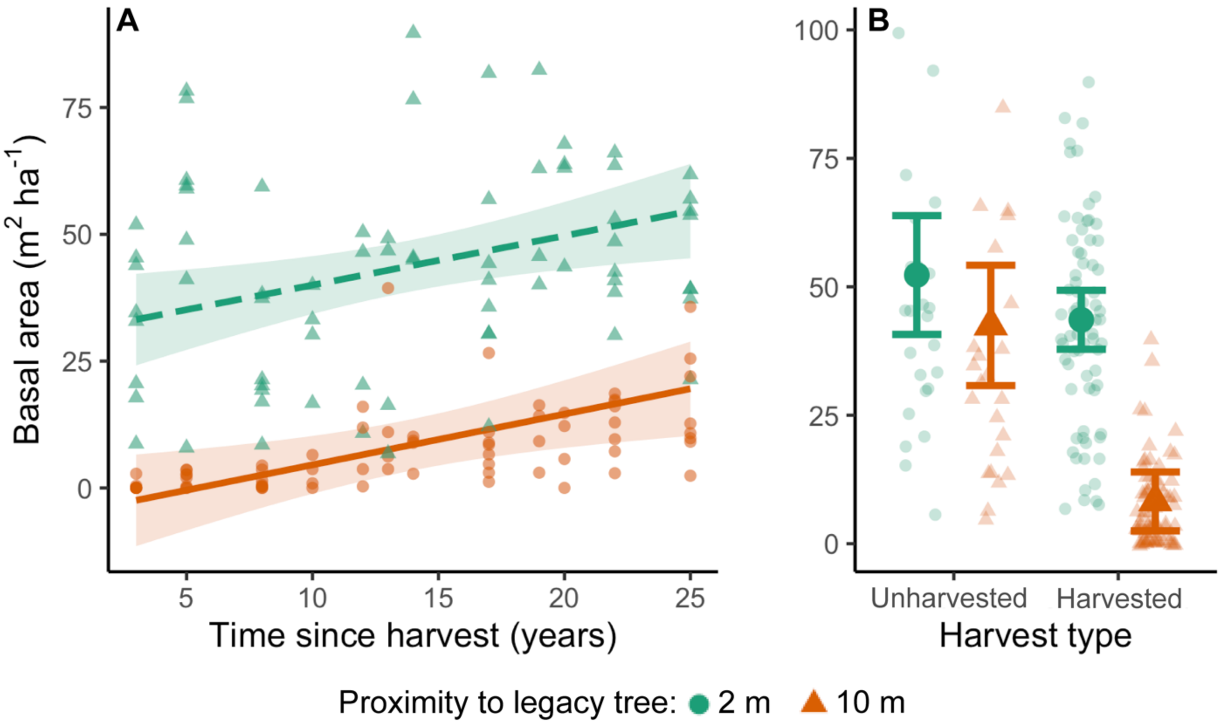


**Fig. S2** Variation in tree basal area in plots 2 m and 10m from legacy overstory trees across the (A) shelterwood chronosequence and (B) in the harvested stands managed through regeneration treatments (*n* = 144 plots) versus the unharvested controls (*n* = 48 plots).

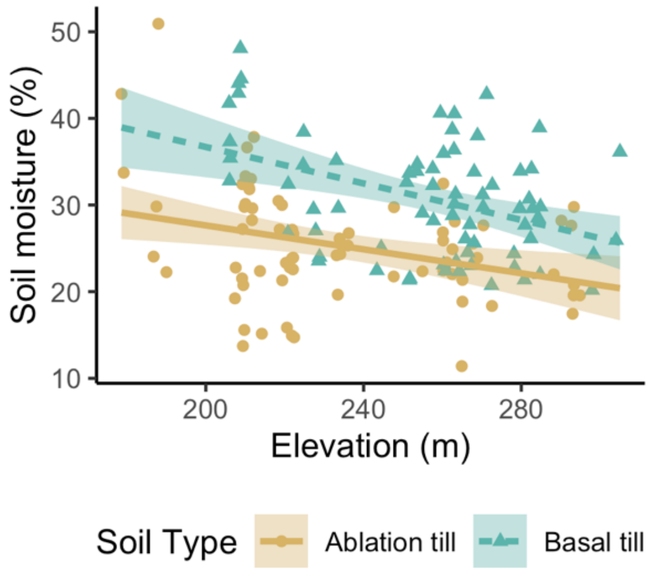


**Fig. S3** Relationship between elevation, soil type, and soil moisture (volumetric water content) in the harvested stands managed to promote regeneration (*n* = 144 plots).

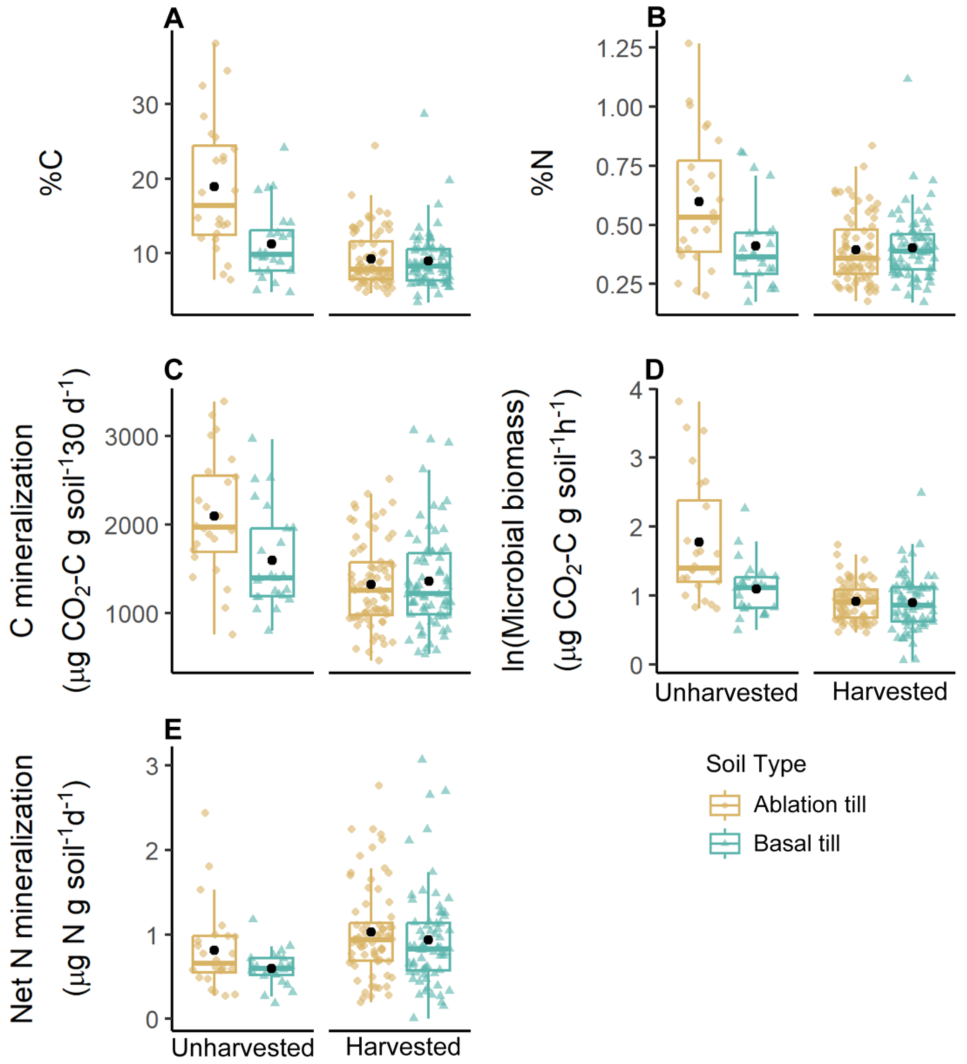


**Fig. S4** The effects of timber harvesting on surface soil carbon (C) and nitrogen (N) content (A, B) and availability (C-E) varied by soil type. Relative to the unharvested stands, the harvested stands managed to promote tree regeneration had more pronounced effects on C and N in the drier and less fertile ablation till soil type compared to the more moist and fertile basal tills (Fig. S3). Black points show the mean values, boxplots show the median, first, and third quartiles, and whiskers show the upper (1.5 × interquartile range) and lower range (1.5 × interquartile range), excluding outliers.

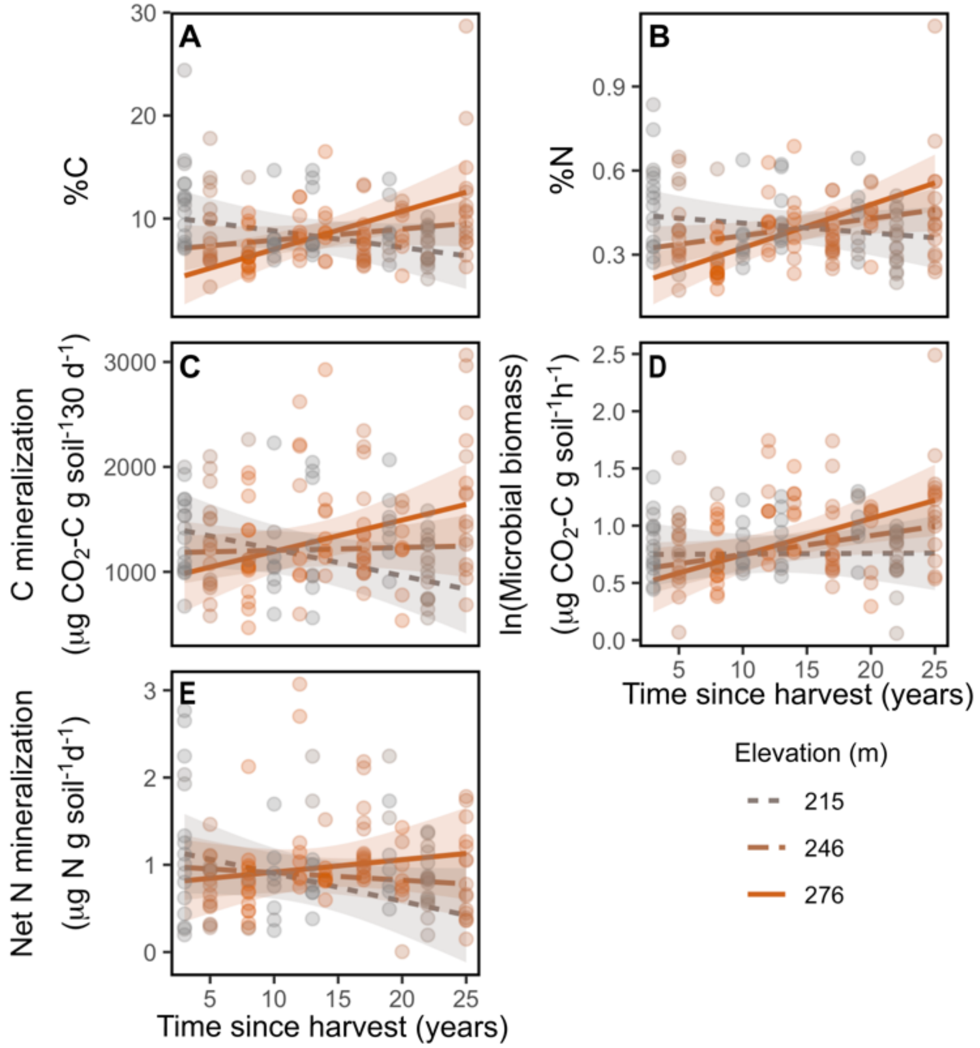


**Fig. S5** The recovery of surface soil carbon(C) and nitrogen (N) content and availability over time since harvest in the 22-year chronosequence (*n* = 144) was contingent on elevation. Carbon and N content and availability increased over time at the higher elevation sites but declined or remained constant at the lower elevation sites. The regression lines depict the modeled effects of the C and N content and availability from the linear mixed effects models (Tables S5-S6), which include plot pair and pair nested within stand as random effects.

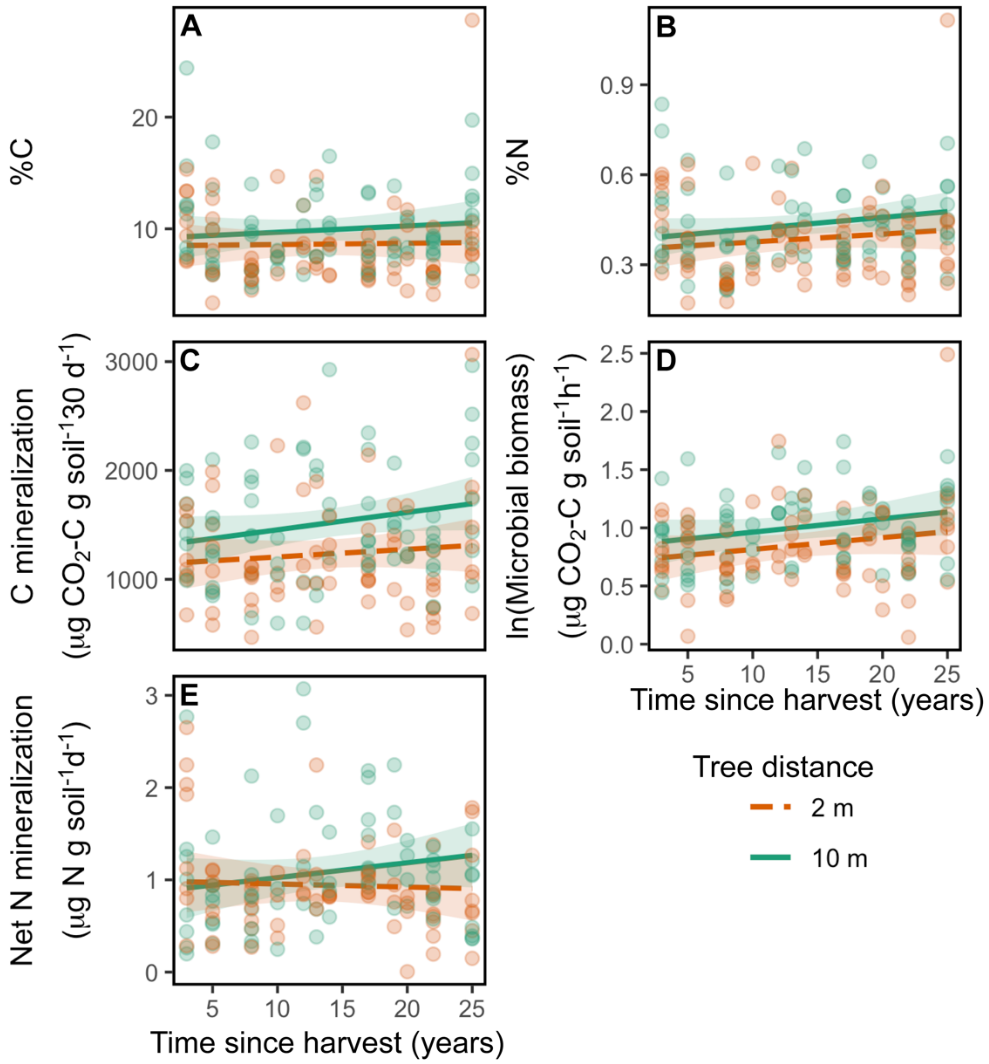


**Fig. S6** Proximity to legacy overstory trees enhanced surface soil C and N content in the harvested stands (*n* = 144 plots), but these effects were largely invariant with time since harvest, with the exception of net N mineralization (E). The regression lines depict the modeled effects of the C and N content and availability from the linear mixed effects models (Tables S5-S6), which include plot pair and pair nested within stand as random effects.

**Table S1** The frequency and abundance of all U.S. soil series in oak-hardwood forest stands at Yale-Myers Forest. Data from Web Soil Survey (NRCS 2019).

| U.S. Soil Types (NRCS, 2019) | Area (Ha) | % of total area | % of stands |
|---|---|---|---|
| Nipmuck-Brookfield complex | 36.2 | 39.3 | 61.1 |
| Paxton and Montauk fine sandy loams | 23.7 | 25.7 | 44.4 |
| Woodbridge fine sandy loams | 19.3 | 20.9 | 38.9 |
| Canton and Charlton fine sandy loams | 8.67 | 9.42 | 22.2 |
| Ridgebury, Leicester, and Whitman soils | 2.83 | 3.08 | 22.2 |
| Nipmuck-Brimfield-rock outcrop complex | 0.958 | 1.04 | 11.1 |
| Sutton fine sandy loams | 0.326 | 0.354 | 11.1 |
| Timakwa and Natchaug soils | 0.161 | 0.175 | 5.56 |

**Table S2** Harvest years of sampled stands across the 22-year chronosequence for each of the two dominant U.S. soil series (NRCS, 2019) used to stratify sampling by variation in geology and parent material.

| U.S. Soil Type (NRCS, 2019) | Harvest Year |
|---|---|
| Nipmuck-Brookfield Complex Series (Ablation tills) | 1994 |
| | 1997 |
| | 2000 |
| | 2002 |
| | 2006 |
| | 2009 |
| | 2011 |
| | 2014 |
| | 2016 |
| Paxton-Montauk and Woodbridge Series (Basal tills) | 1994 |
| | 1997 |
| | 1999 |
| | 2002 |
| | 2005 |
| | 2007 |
| | 2011 |
| | 2014 |
| | 2016 |

**Table S3** Standard deviation (SD) and proportion of variance explained for the 15 principal components (PCs) of soil variables for the PCA to compare the harvested ($n$ = 144) and unharvested ($n$ = 48) plots (Fig. 2a) and for the PCA of the time since harvest chronosequence (Fig. 2b).

| PC | Harvested vs. unharvested | | Time since harvest chronosequence | |
|---|---|---|---|---|
| | SD | Proportion of variance | SD | Proportion of variance |
| 1 | 2.75 | 0.503 | 2.67 | 0.475 |
| 2 | 1.28 | 0.110 | 1.33 | 0.118 |
| 3 | 1.17 | 0.091 | 1.15 | 0.088 |
| 4 | 1.08 | 0.078 | 1.07 | 0.077 |
| 5 | 0.93 | 0.058 | 0.94 | 0.059 |
| 6 | 0.81 | 0.044 | 0.87 | 0.050 |
| 7 | 0.70 | 0.033 | 0.71 | 0.033 |
| 8 | 0.55 | 0.020 | 0.59 | 0.023 |
| 9 | 0.54 | 0.019 | 0.55 | 0.020 |
| 10 | 0.43 | 0.012 | 0.53 | 0.019 |
| 11 | 0.41 | 0.011 | 0.46 | 0.014 |
| 12 | 0.36 | 0.009 | 0.40 | 0.011 |
| 13 | 0.29 | 0.005 | 0.33 | 0.007 |
| 14 | 0.25 | 0.004 | 0.25 | 0.004 |
| 15 | 0.14 | 0.001 | 0.16 | 0.002 |

**Table S4** Loading factors for the 15 principal components (PCs) of soil variables for the PCA comparing the harvested ($n$ = 144) to unharvested ($n$ = 48) plots (Fig 2a).

| PC | Loadings | | | | | | | | | | | | | | |
|---|---|---|---|---|---|---|---|---|---|---|---|---|---|---|---|
| | VWC | FWM | CWM | BD | ph | GWC | WHC | SOM | N Min | SIR | Cmin | N | C | Oa | Soil temp |
| 1 | 0.09 | -0.04 | -0.04 | 0.32 | 0.26 | -0.30 | -0.34 | -0.35 | -0.07 | -0.32 | -0.33 | -0.33 | -0.35 | -0.23 | 0.11 |
| 2 | 0.58 | 0.40 | 0.25 | 0.17 | 0.17 | 0.31 | 0.06 | 0.06 | 0.28 | -0.09 | -0.03 | 0.21 | 0.03 | -0.38 | -0.10 |
| 3 | 0.04 | 0.32 | 0.42 | 0.07 | -0.33 | -0.01 | -0.10 | -0.03 | -0.52 | -0.12 | -0.13 | -0.12 | 0.00 | 0.25 | -0.47 |
| 4 | 0.44 | -0.46 | -0.52 | 0.04 | 0.03 | 0.23 | 0.08 | 0.03 | -0.21 | -0.11 | -0.04 | -0.01 | -0.05 | 0.09 | -0.45 |
| 5 | 0.30 | -0.41 | 0.45 | -0.03 | 0.21 | 0.17 | -0.02 | 0.06 | -0.44 | 0.06 | 0.02 | 0.06 | 0.05 | 0.11 | 0.50 |
| 6 | 0.16 | -0.42 | 0.39 | 0.08 | -0.36 | -0.05 | 0.00 | -0.11 | 0.61 | -0.04 | -0.08 | -0.15 | -0.07 | 0.29 | -0.11 |
| 7 | 0.26 | 0.37 | -0.31 | -0.06 | -0.27 | 0.18 | 0.07 | 0.02 | 0.05 | -0.29 | -0.04 | -0.16 | -0.14 | 0.47 | 0.48 |
| 8 | 0.10 | 0.08 | -0.17 | 0.61 | 0.00 | -0.18 | -0.18 | -0.06 | 0.01 | 0.41 | -0.20 | 0.26 | 0.38 | 0.30 | 0.08 |
| 9 | 0.01 | -0.16 | -0.10 | 0.06 | -0.69 | -0.01 | 0.00 | 0.04 | -0.15 | -0.12 | -0.24 | 0.14 | 0.19 | -0.54 | 0.23 |
| 10 | 0.30 | 0.09 | -0.08 | -0.20 | -0.22 | 0.01 | -0.33 | -0.25 | -0.04 | 0.62 | 0.31 | -0.31 | -0.17 | -0.16 | 0.03 |
| 11 | 0.03 | 0.02 | -0.03 | -0.53 | 0.09 | 0.08 | -0.52 | 0.14 | 0.12 | 0.06 | -0.57 | 0.22 | 0.07 | 0.10 | -0.05 |
| 12 | 0.14 | -0.03 | -0.02 | -0.12 | -0.04 | -0.26 | -0.42 | -0.21 | 0.01 | -0.43 | 0.54 | 0.32 | 0.29 | 0.06 | -0.03 |
| 13 | 0.05 | 0.05 | 0.01 | -0.28 | 0.05 | -0.02 | 0.42 | -0.79 | -0.04 | 0.05 | -0.22 | 0.18 | 0.17 | 0.04 | 0.00 |
| 14 | 0.39 | 0.05 | 0.00 | -0.24 | 0.04 | -0.76 | 0.31 | 0.32 | -0.03 | 0.06 | -0.09 | -0.06 | 0.05 | -0.01 | -0.01 |
| 15 | 0.01 | 0.01 | -0.01 | -0.07 | 0.13 | 0.12 | -0.03 | 0.02 | 0.06 | -0.11 | -0.05 | -0.64 | 0.73 | -0.07 | -0.01 |

Abbreviations are as follows: volumetric water content (VWC), fine woody material (FWM), coarse woody material (CWM), bulk density (BD), gravimetric water content (GWC), water holding capacity (WHC), soil organic matter content (SOM), net potential N mineralization (N Min), microbial biomass as estimated with substrate induced respiration (SIR), carbon mineralization (Cmin), carbon content (C), nitrogen content (N), Oa horizon depth (Oa), soil temperature (Soil temp).

**Table S5** Loading factors for the 15 principal components (PCs) of soil variables for the PCA for the plots in the 22-year time since harvest chronosequence (*n* = 144) (Fig. 2b).

| PC | Loadings | | | | | | | | | | | | | | |
|---|---|---|---|---|---|---|---|---|---|---|---|---|---|---|---|
| | VWC | FWM | CWM | BD | ph | GWC | WHC | SOM | N Min | SIR | Cmin | N | C | Oa | Soil temp |
| 1 | -0.04 | -0.04 | -0.01 | 0.32 | 0.22 | -0.34 | -0.34 | -0.35 | -0.09 | -0.29 | -0.33 | -0.34 | -0.35 | -0.21 | 0.11 |
| 2 | 0.50 | 0.36 | 0.35 | 0.21 | -0.30 | 0.18 | -0.04 | 0.00 | -0.23 | -0.26 | -0.18 | 0.01 | 0.05 | 0.00 | -0.44 |
| 3 | 0.11 | -0.57 | -0.39 | 0.01 | -0.08 | 0.06 | 0.01 | -0.03 | -0.46 | -0.02 | -0.04 | -0.15 | -0.05 | 0.40 | -0.32 |
| 4 | 0.53 | -0.17 | -0.48 | 0.08 | 0.24 | 0.18 | 0.05 | 0.06 | 0.31 | -0.08 | -0.07 | 0.20 | 0.03 | -0.45 | -0.04 |
| 5 | 0.28 | -0.14 | 0.34 | -0.08 | 0.44 | 0.14 | -0.08 | 0.12 | -0.55 | 0.07 | 0.03 | 0.15 | 0.10 | -0.05 | 0.46 |
| 6 | 0.33 | -0.49 | 0.38 | 0.06 | -0.33 | 0.01 | -0.03 | -0.09 | 0.46 | 0.00 | -0.09 | -0.09 | -0.03 | 0.25 | 0.32 |
| 7 | 0.08 | 0.36 | -0.44 | 0.08 | -0.24 | 0.08 | 0.08 | 0.03 | -0.09 | -0.35 | -0.12 | 0.04 | 0.07 | 0.34 | 0.58 |
| 8 | 0.28 | 0.30 | -0.05 | 0.14 | 0.51 | -0.05 | -0.13 | -0.06 | 0.25 | 0.29 | 0.12 | -0.12 | -0.06 | 0.57 | -0.13 |
| 9 | 0.26 | 0.19 | -0.15 | -0.41 | -0.30 | 0.16 | -0.22 | -0.19 | -0.12 | 0.41 | 0.17 | -0.33 | -0.35 | -0.20 | 0.13 |
| 10 | 0.04 | 0.02 | -0.11 | 0.63 | -0.26 | -0.24 | 0.00 | -0.01 | -0.16 | 0.59 | -0.01 | 0.09 | 0.23 | -0.14 | 0.12 |
| 11 | 0.12 | -0.03 | 0.05 | 0.23 | 0.01 | -0.10 | 0.32 | -0.20 | -0.06 | -0.28 | 0.80 | -0.19 | -0.05 | -0.10 | 0.04 |
| 12 | 0.05 | -0.04 | -0.09 | -0.14 | -0.12 | -0.23 | -0.72 | -0.16 | 0.01 | -0.16 | 0.32 | 0.36 | 0.32 | 0.04 | -0.05 |
| 13 | 0.09 | 0.04 | 0.01 | -0.25 | 0.06 | -0.16 | 0.38 | -0.77 | -0.06 | 0.09 | -0.21 | 0.25 | 0.21 | 0.03 | -0.02 |
| 14 | 0.30 | 0.04 | -0.02 | -0.32 | -0.04 | -0.77 | 0.18 | 0.38 | -0.04 | -0.03 | -0.07 | -0.16 | 0.08 | -0.02 | -0.02 |
| 15 | 0.04 | 0.00 | 0.02 | 0.06 | -0.09 | -0.18 | 0.08 | 0.06 | -0.06 | 0.03 | 0.07 | 0.64 | -0.72 | 0.11 | -0.01 |

Abbreviations are as follows: volumetric water content (VWC), fine woody material (FWM), coarse woody material (CWM), bulk density (BD), gravimetric water content (GWC), water holding capacity (WHC), soil organic matter content (SOM), net potential N mineralization (N Min), microbial biomass as estimated with substrate induced respiration (SIR), carbon mineralization (Cmin), carbon content (C), nitrogen content (N), Oa horizon depth (Oa), soil temperature (Soil temp).

**Table S3** Unstandardized coefficients of models assessing the effects of timber harvesting on surface soil carbon (C) and nitrogen (N) concentrations and C:N in the harvested stands versus unharvested controls (Q1).

| Predictors | Unstandardized coefficients (SE) | | |
|---|---|---|---|
| | %C | %N | C:N[1] |
| Intercept | **13.0 (2.0)*** ** | **0.48 (0.070)*** ** | **3.29 (0.052)*** ** |
| Harvest | **-0.5.06 (2.14)*** | -0.11 (0.075) | **-0.23 (0.059)*** ** |
| Tree | -0.69 (1.78) | -0.029 (0.059) | -0.0033 (0.018) |
| Soil Type[2] | **6.40 (2.16)** ** | **0.15 (0.074)*** | **0.13 (0.051)*** |
| Harvest × Tree | 2.01 (1.84) | 0.079 (0.062) | **0.016 (0.022)** |
| Harvest × Soil Type | **-5.71 (2.28)*** | **-0.15 (0.079)** | -0.054 (0.059) |
| $R^2$ Fixed /Full | 0.45/0.69 | 0.21/0.52 | 0.43/0.82 |

[1]Variable is natural log-transformed
[2]Soil type is a binary variable, where ablation till soils = 1 and basal till soils = 0
Significance is denoted in bold and with asterisks as follows: $P < 0.01$ is bold only, $P < 0.05$ *, $P < 0.01$ **, $P < 0.001$ ***

**Table S4** Unstandardized coefficients of models assessing the effects of timber harvesting on carbon (C) and nitrogen (N) availability (C mineralization, microbial biomass, and N mineralization) in the harvested stands versus unharvested controls (Q1).

| Predictors | Unstandardized Coefficients (SE) | | |
|---|---|---|---|
| | C mineralization (ug $CO_2$-C g soil$^{-1}$ 30 d$^{-1}$) | Microbial biomass[1] (ug $CO_2$-C g soil$^{-1}$ h$^{-1}$) | Net N mineralization (ug N g soil$^{-1}$ d$^{-1}$) |
| Intercept | **1696.1 (193.6)*** ** | **1.22 (0.20)*** ** | **0.65 (0.15)*** ** |
| Harvest | **-497.5 (212.6)*** | **-0.42 (0.21)** | 0.21 (0.18) |
| Tree | -77.46 (166.9) | -0.042 (0.18) | 0.094 (0.10) |
| Soil Type[2] | **454.2 (211.7)*** | **0.58 (0.21)**** | 0.13 (0.14) |
| Harvest × Tree | **364.7 (181.3)*** | 0.20 (0.19) | 0.050 (0.13) |
| Harvest × Soil Type | **-452.1 (233.1)** | **-0.51 (0.22)*** | -0.028 (0.17) |
| $R^2$ Fixed /Full | 0.25/0.47 | 0.42/0.64 | 0.049/0.21 |

[1]Variable is natural log-transformed
[2]Soil type is a binary variable, where ablation till soils = 1 and basal till soils = 0
Significance is denoted in bold and with asterisks as follows: $P < 0.01$ is bold only, $P < 0.05$ *, $P < 0.01$ **, $P < 0.001$ ***

**Table S5** Unstandardized coefficients of models assessing the effects of time since harvest on the recovery of surface soil carbon (C) and nitrogen (N) concentrations and C:N in the 22-year chronosequence (Q2).

| Predictors | Unstandardized coefficients (SE) | | |
|---|---|---|---|
| | %C | %N | C:N[1] |
| Intercept | **37.6 (8.4)**** | **1.5 (0.3)***** | **3.4 (0.4)***** |
| Years | **-2.23 (0.64)**** | **-0.078 (0.021)**** | -0.044 (0.030) |
| Tree | 0.65 (0.96) | 0.032 (0.038) | -0.040 (0.027) |
| Soil Type[2] | **3.19 (1.32)*** | 0.082 (0.049) | 0.095 (0.058) |
| Elevation (m) | **-0.12 (0.03)**** | **-0.0045 (0.0011)**** | -0.0010 (0.0016) |
| FWM (pieces $m^{-2}$) | **-1.06 (0.29)***** | **-0.033 (0.011)**** | -0.015 (0.010) |
| Years × Tree | 0.04 (0.06) | 0.0012 (0.0025) | **0.0038 (0.0018)*** |
| Years × Soil Type | **-0.18 (0.08)*** | **-0.0067 (0.0031)*** | -0.00023 (0.0035) |
| Years × Elevation | **0.009 (0.003)**** | **0.00031 (0.00008)**** | 0.00015 (0.00012) |
| Years × FWM | **0.11 (0.02)***** | **0.0040 (0.00087)***** | 0.00084 (0.00075) |
| $R^2$ Fixed /Full | 0.29/0.53 | 0.30/0.47 | 0.17/0.73 |

[1]Variable is natural log-transformed
[2]Soil type is a binary variable, where ablation till soils = 1 and basal till soils = 0
Significance is denoted in bold and with asterisks as follows: $P < 0.01$ is bold only, $P < 0.05$ *, $P < 0.01$ **, $P < 0.001$ ***

**Table S6** Unstandardized coefficients of models assessing the effects of time since harvest on carbon (C) and nitrogen (N) availability (C mineralization, microbial biomass, and N mineralization) in the 22-year chronosequence (Q2).

| Predictors | Unstandardized Coefficients (SE) | | |
|---|---|---|---|
| | C mineralization (ug $CO_2$-C g soil$^{-1}$ 30 d$^{-1}$) | Microbial biomass[1] (ug $CO_2$-C g soil$^{-1}$ h$^{-1}$) | Net N mineralization (ug N g soil$^{-1}$ d$^{-1}$) |
| Intercept | **3635.8 (1049.2)*** | **2.00 (0.85)*** | **2.93 (1.43)** |
| Years | **-238.3 (79.7)*** | **-0.12 (0.065)** | **-0.21 (0.11)** |
| Tree | 159.9 (147.6) | 0.13 (0.099) | -0.13 (0.18) |
| Soil Type[2] | -93.1 (198.4) | **0.25 (0.13)** | 0.021 (0.20) |
| Elevation | **-9.38 (4.15)** | -0.0052 (0.0034) | -0.0074 (0.0057) |
| FWM (pieces m$^{-2}$) | **-81.0 (44.4)** | **-0.067 (0.029)*** | -0.068 (0.049) |
| Years × Tree | **8.96 (9.47)** | 0.0014 (0.0063) | **0.019 (0.011)** |
| Years × Soil Type | 9.47 (12.5) | -0.012 (0.0078) | 0.011 (0.012) |
| Years × Elevation | **0.90 (0.32)*** | **0.00051 (0.00026)** | 0.00076 (0.00044) |
| Years × FWM | **9.94 (3.44)**** | **0.0067 (0.0022)**** | **0.011 (0.0038)**** |
| $R^2$ Fixed /Full | 0.19/0.42 | 0.19/0.43 | 0.13/0.31 |

[1]Variable is natural log-transformed
[2]Soil type is a binary variable, where ablation till soils = 1 and basal till soils = 0
Significance is denoted in bold and with asterisks as follows: $P < 0.01$ is bold only, $P < 0.05$ *, $P < 0.01$ **, $P < 0.001$ ***
